# Bayesian Parameter Estimation for Dynamical Models in Systems Biology

**DOI:** 10.1101/2022.04.11.487931

**Authors:** Nathaniel J. Linden, Boris Kramer, Padmini Rangamani

## Abstract

Dynamical systems modeling, particularly via systems of ordinary differential equations, has been used to effectively capture the temporal behavior of different biochemical components in signal transduction networks. Despite the recent advances in experimental measurements, including sensor development and ‘-omics’ studies that have helped populate protein-protein interaction networks in great detail, systems biology modeling lacks systematic methods to estimate kinetic parameters and quantify associated uncertainties. This is because of multiple reasons, including sparse and noisy experimental measurements, lack of detailed molecular mechanisms underlying the reactions, and missing biochemical interactions. Additionally, the inherent nonlinearities with respect to the states and parameters associated with the system of differential equations further compound the challenges of parameter estimation. In this study, we propose a comprehensive framework for Bayesian parameter estimation and complete quantification of the effects of uncertainties in the data and models. We apply these methods to a series of signaling models of increasing mathematical complexity. Systematic analysis of these dynamical systems showed that parameter estimation depends on data sparsity, noise level, and model structure, including the existence of multiple steady states. These results highlight how focused uncertainty quantification can enrich systems biology modeling and enable additional quantitative analyses for parameter estimation.

## 1. Introduction

Mathematical modeling is an integral part of systems biology; indeed, the use of approaches from dynamical systems analyses resulted in a paradigmatic shift in our understanding of biochemical signal transduction and enabled the identification of the emergent properties of a signaling network [1, 2, 3, 4]. Additionally, mathematical models allow us to investigate the dynamics of biological systems beyond what is experimentally possible [5, 6, 7, 8]. A classical approach to modeling the dynamics of signal transduction is the use of systems of ordinary differential equations (ODEs) [9, 10]. Often these equations include nonlinear functions to capture complex biochemical interactions using MichaelisMenten kinetics and Hill functions for cooperative binding [11]. One of the ongoing challenges in developing and constraining predictive models of signal transduction has been the estimation and identification of the kinetic parameters associated with these reactions and quantifying the associated uncertainty [12, 13, 14]. The use of rigorous, quantitative approaches to estimate kinetic parameters and their uncertainties is in its early stages in systems biology [15, 12, 16, 17, 18] even though such methods are far more prevalent in the greater computational science community under the field of uncertainty quantification (UQ) [19, 20, 21, 22].

There are many sources of uncertainty in dynamical systems modeling of signal transduction, including the model structure itself, the values of model parameters, and the quality of the data used for model calibration. Uncertainty in the model equations, known as model form or topological uncertainty [23, 24, 25, 26] often arises during model development. However, the reaction fluxes for many biochemical reactions (ODE formulations) can be established in terms of classical rate equations [9, 10, 11]. The more significant challenge is establishing suitable model parameters, including the kinetic rate constants, for various flux terms [16]. Direct measurement of these parameters often occurs in isolated reaction systems and does not capture the complexity of the large network of reactions represented by dynamical systems models. Estimating these biological parameters and identifying any remaining uncertainties requires selecting a statistical model and then learning the distribution of these parameters from available data [19, 27]. From this viewpoint the biological parameters are random variables that either have parametric or nonparametric distributions. However, parameter estimation is complicated by the noisy, sparse (few time points), and incomplete nature of data found in systems biology (few or select readouts due to experimental limitations) that introduce uncertainties in the biological parameters [28, 29, 14, 30, 31]. In the face of these complicating factors, there is a need for statistical modeling of parameters that enables uncertainty quantification.

A comprehensive parameter estimation and UQ framework should consider the impact of structural parameter identifiability and parameter sensitivity [16, 29, 32, 33, 34]. Structural parameter identifiability analysis reveals which of the parameters can be estimated given a specific dynamical systems model and a set of measurable outputs [28, 29, 35, 36]. A parameter is globally structurally identifiable if there is only one unique model output for each value of that parameter [29]. Parameters that do not meet this criterion are deemed structurally nonidentifiable and cannot successfully be estimated from the specified model outputs. Structural nonidentifiabilities can arise due to complex nonlinear equations and incomplete experimental data that only measures a subset of the system’s states. Additionally, parametric sensitivity analysis [19, 37, 38] quantifies how sensitive a model output is to variations in the model parameters. Gutenkunst et al. [28] found that most models in systems biology contain parameters with a wide range of sensitivities, which they termed ‘sloppy’. Despite this challenge, sensitivity analysis helps rank the set of identifiable parameters by their contributions to specified model outputs, [39] as was done in Mortlock et al. [40] for the prolactin-mediated JAK-STAT signaling pathway. This analysis enabled them to select a subset of 33 out of 60 total parameters that significantly contribute to variations in the model outputs.

Commonly used methods to estimate parameters for systems biology models include frequentist [17] and Bayesian approaches [18]. In the frequentist setting, parameter estimation is formulated as an optimization problem, and the solution to the parameter estimation problem is the set of parameters that best recapitulates the data [19]. Additionally, frequentist approaches quantify uncertainty via estimated confidence intervals around the optimal parameters [41, 42, 19]. In contrast, in the Bayesian perspective, parameters are assumed to be random variables whose unknown probability distributions, called *posterior distributions*, quantify the probability of assuming any value in the parameter space [19, 21, 27]. The advantage of Bayesian approaches comes from their ability to characterize the entire posterior distribution and quantify the uncertainty in parameter estimates via credible intervals [27]. Many methods have been developed for Bayesian parameter estimation [18, 43, 44, 45, 46, 47, 48, 49] that all aim to characterize the posterior distribution, by leveraging Bayes’ rule [19, 27]. For example, Mortlock et al. [40] successfully used Bayesian estimation to study the uncertainty in the model predictions and assess the statistical significance of their modeling results.

Despite the successes of Bayesian parameter estimation in systems biology [12, 15, 16, 18], failure to account for all sources of uncertainty in a model can significantly inhibit parameter estimation and uncertainty quantification [16, 26, 28]. Thus, a comprehensive framework for UQ in systems biology should include rigorous accounting of uncertainties in the model structure, nonidentifiable parameters, mixed parameter sensitivities, and noisy, sparse, or incomplete experimental data. While identifiability and sensitivity analyses are typically performed prior to parameter estimation [16, 29], accounting for model form uncertainty requires us to consider a stochastic model instead of a deterministic one [23, 26, 50, 30]. One promising approach to account for model form uncertainty is the Unscented Kalman filter Markov chain Monte Carlo (UKF-MCMC) method [26, 51]. This method includes statistical models for noisy data and model form uncertainty simultaneously; however, it has not been adapted for dealing with the unique challenges in system biology. Similarly, the parameter estimation and model selection method in [30] accounts for model form uncertainty with extended Kalman filtering but does not provide complete uncertainty estimates because it takes a frequentist approach for parameter estimation. Thus, there is a need for a framework that combines structural identifiability analysis, global sensitivity analysis, a statistical model for data and model form uncertainty, and Bayesian parameter estimation for comprehensive UQ of dynamical models in systems biology.

In this work, we propose a comprehensive workflow that addresses limitations of Bayesian parameter estimation due to parameter nonidentifiabilities and accounts for uncertainties in the data and the model structure by building on the Bayesian framework and the UKF-MCMC method. This workflow begins with structural identifiability analysis [36, 52] and global sensitivity analysis [37, 53] and then extends the UKF-MCMC [26, 51] approach for systems biology by leveraging the constrained interval unscented Kalman filter (CIUKF) [54]. We applied this framework to three systems biology models of increasing complexity, including a simple two-state model [55], a model of the core mitogen-activated protein kinase (MAPK) signaling pathway [56], and a phenomenological model of synaptic plasticity to capture long-term potentiation/depression [57]. We found that even in simple models, estimation of parameters depends on the level of data noise and data sparsity. Finally, the framework enabled uncertainty quantification for model structures that include non-linearities and multistability. In all of these cases, we leveraged identifiability and sensitivity analyses to narrow the subset of parameters for estimation and then used Bayesian estimation to determine the role of model structure in parameter estimation. These results establish an uncertainty quantification-focused approach to systems biology that can enable rigorous parameter estimation and analysis.

## 2. Methods

This section describes the technical details of the comprehensive framework for uncertainty quantification (see Fig 1) proposed in section 2.1. Next, section 2.2 introduces dynamical systems biology models and the parameter estimation problem. Sections 2.3 and 2.4 overview structural identifiability and global sensitivity analyses, respectively, to reduce the dimension of the parameter space. Then section 2.5 introduces Bayesian estimation, section 2.6 outlines the CIUKF-MCMC algorithm, and section 2.7 describes the constrained interval unscented Kalman filter in detail. Following this, section 2.8 discusses how to construct prior distributions and section 2.9 details Markov chain Monte Carlo sampling. Lastly, section 2.10 discusses output uncertainty propagation with ensemble modeling, section 2.11 highlights choosing point estimators for the parameters, section 2.12 delineates synthetic data generation and section 2.13 outlines limit cycle analysis.

**Figure 1:**
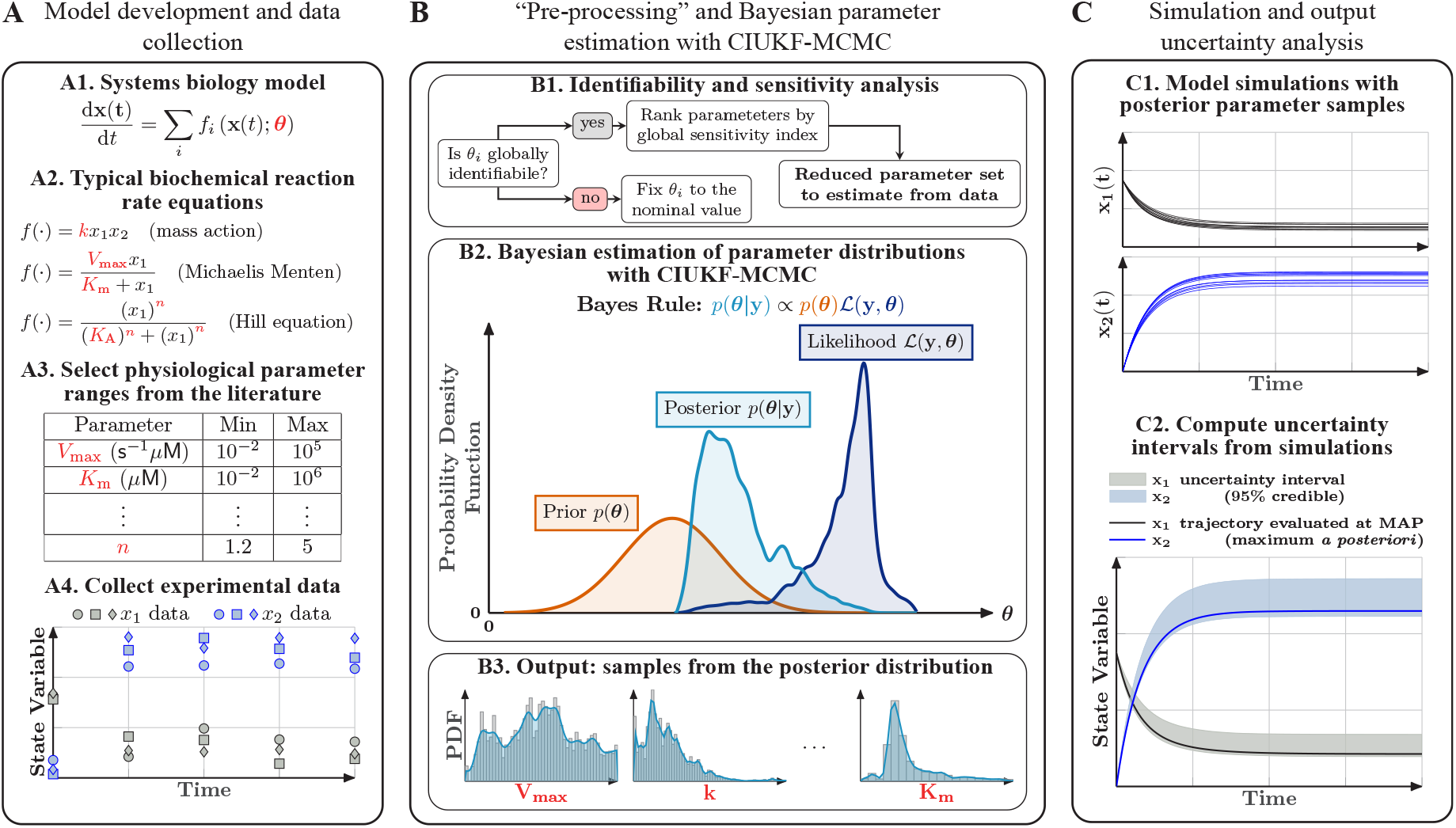
A comprehensive Bayesian parameter estimation and uncertainty quantification framework for dynamical models in systems biology. (**A**) Model development in systems biology begins with model construction and data collection. Dynamical models in systems biology typically involve a system of ODEs that capture the dynamics of the concentrations of different chemical species in the system (A1). The reaction rates associated with these concentration changes are usually mass action, Michaelis Menten kinetics, or cooperative kinetics represented by the Hill equation (A2). The free parameters in these models include kinetic rate constants, e.g. *k, V*_max_, equilibrium constants, e.g. *K*_*m*_, *K*_*A*_, and Hill coefficients, e.g. *n*. These parameters are first constrained by *best guess* values based on physiological ranges and typical values of model parameters from the literature (A3). Finally, the model needs experimental data for validation; this data can either be from published work or new experiments. (**B**) Parameter preprocessing and Bayesian parameter estimation with the CIUKF-MCMC algorithm. First structural identifiability and global sensitivity analyses on the entire parameter set reduce the set of free parameters that can be estimated (B1). Next, we perform Bayesian parameter estimation for this reduced set of parameters to learn their posterior distributions. The posterior distribution is the parameter distribution conditioned on the data (B2). Bayes’ rule relates the posterior distribution to the product of the prior distribution and the likelihood function. The prior distribution encodes known information about the parameters and the likelihood function (which requires simulating the model) measures the misfit between predictions and the data. A state-constrained Unscented Kalman filter approximates the likelihood function to account for uncertainty in the model equations. Although Bayes’ rule provides a means to evaluate the posterior distribution at specific points in the parameter space, we use Markov chain Monte Carlo (MCMC) sampling (B3) to characterize the entire distribution. (**C**) The posterior distributions enable uncertainty analysis of model outputs through ensemble simulation. We perform simulations using the posterior parameter samples to propagate parameter uncertainty through the model (C1). Statistical analysis of the ensemble enables us to compute uncertainty intervals and study various system behaviors, for example, the statistics of the steady state values (C2).

### 2.1 A framework for comprehensive uncertainty quantification for dynamical models in systems biology

This section previews the proposed comprehensive framework for parameter estimation and uncertainty quantification of dynamical models in systems biology. Figure 1 outlines the framework and its components, which are then described in much more detail in the subsequent sections. The proposed framework follows three main steps. First, we assume that dynamical models of intracellular signal transduction (Fig 1.A1) use classical biochemical rate laws, such as mass action kinetics, Michaelis-Menten kinetics, and Hill functions [9, 10, 11] (Fig 1.A2). The key challenge to applying these models is estimating the associated parameters, such as the rate constants *k* and *V*_max_, equilibrium coefficients *K*_*m*_ and *K*_*A*_, and Hill coefficients *n* in Fig 1.A2, from available experimental data (Fig 1.A4). The comprehensive framework uses Bayesian inference to estimate a statistical model (a probability distribution; see Fig 1.B) for the model parameters given a set of noisy measurement data and a specific model form.

We argue that identifiability and sensitivity analysis are necessary steps to perform before parameter estimation (Fig 1.B1). To eliminate uncertainty due to nonidentifiable parameters, we perform global structural identifiability analysis using the Structural Identifiability Analyzer (SIAN) [36, 52] (see section 2.3 for details). The nonidentifiable parameters are fixed to their nominal values from the literature or based on their physiological ranges. Next, variance-based global sensitivity analysis [19, 37] is performed to rank the identifiable parameters in order of their contributions to the variance of the model outputs (see section 2.4 for details). A subset of the identifiable parameters with the largest sensitivity indices is selected for parameter estimation. The remaining model parameters are fixed to their nominal values in the same fashion as nonidentifiable parameters.

Bayesian parameter estimation completely characterizes uncertainty in the model parameters by estimating a nonparametric statistical model. Bayes’ rule (see Fig 1.B2) provides the *best guess* distribution, called the *posterior distribution*, for the parameters starting from an initial guess (the prior distribution) that is transformed by the available experimental data with the likelihood function. The likelihood function measures the mismatch between the data and the model predictions and returns higher probabilities for parameter sets that produce outputs *closer* to the data. We use the CIUKF-MCMC algorithm [26, 51] to approximate the likelihood function and account for uncertainty in the model formulation, data, and parameters. Markov chain Monte Carlo sampling, with either delayed rejection adaptive Metropolis [58] or the affine invariant ensemble sampler [59], generates a set of samples that represents the posterior distribution (Fig 1.B3).

Third, we leverage the posterior distribution to quantify how uncertainty in the model parameters affects uncertainty in the model predictions (Fig 1.C). An ensemble simulations with the parameter samples generates sets of trajectories (see Fig 1.C1) that capture the uncertainty in the predicted dynamics. Computing uncertainty intervals such as the 95% credible intervals presented in Fig 1.C2 provides a visualization of this uncertainty. Notably, credible intervals are different from confidence intervals because credible intervals capture a specified percentage of the samples whereas confidence intervals are random variables that capture regions where estimators will lie at a specified probability level [27] (see Fig 2 for an example of a credible interval). Additionally, statistical analysis of the ensemble enables quantitative analysis of computational modeling results in the same way that running multiple experimental trials enables analysis of experimental results. Before discussing the methods that enable comprehensive uncertainty quantification in the following sections, the next section formally introduces parameter estimation for dynamical models in systems biology.

**Figure 2:**
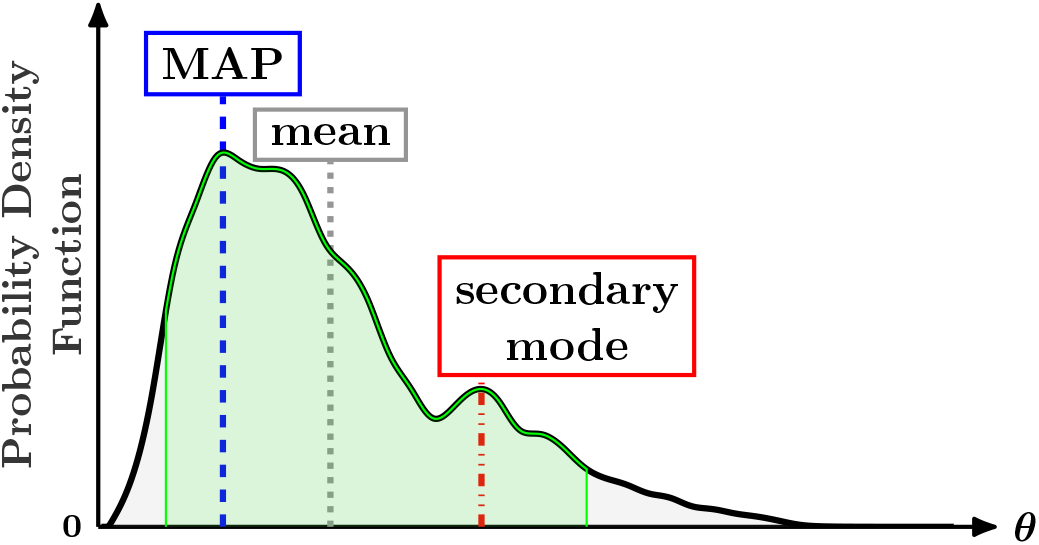
Examples of point estimates depicted on an arbitrary probability density function (black line). The MAP (maximum *a posteriori*) point is located at the most probable point (blue dashed line). The long tail of the distribution shifts the mean (gray dotted line) away from the MAP point. Secondary modes (red dashed-dotted line) can effect the quality of a point estimate. Additionally, the green shaded region highlights the 95% credible interval, the region between the 2.5th and 97.5th percentiles, that is used to capture the uncertainty in an estimate.

### 2.2. Parameter estimation for systems biology models in the form of partially-observed systems of ordinary differential equations

We consider nonlinear ordinary differential equation models of the form

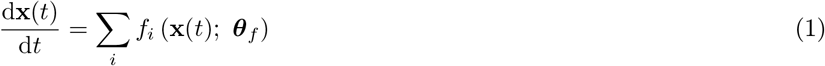

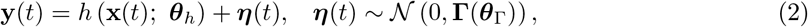

where 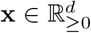 is the state vector of nonnegative species concentrations and **y** ∈ ℝ^*m*^ is the vector of potentially incomplete, *m ≤ d*, measurements of **x**. The functions *f*_*i*_(·; ·) : ℝ^*d*^ *×*ℝ^*p*^ *→*ℝ^*d*^ govern the rates of the involved biochemical reactions and are derived using biochemical theory (see Fig 1.A2 for example terms). Further, 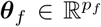 is the vector of biological model parameters, including but not limited to rate constants, binding coefficients, and equilibrium coefficients. The function *h*(·; ·) : ℝ^*d*^ *×*ℝ^*p*^*→* ℝ^*m*^ is the measurement function that maps from the states to the set of observables (experimental data), where ***θ***_*h*_ is the vector of associated parameters. Lastly, the measurements **y**(*t*) are corrupted by independently and identically distributed (iid) Gaussian measurement noise ***η***(*t*) ∈ ℝ^*m*^ with zero mean and covariance matrix **Γ** ∈ ℝ^*m×m*^. The covariance matrix is parameterized by ***θ***_Γ_ ∈ ℝ^*m*^ such that **Γ** (***θ***_Γ_) = diag (***θ***_Γ_). The parameter space is then defined as the multidimensional space of all possible values of ***θ*** = [***θ***_*f*_, ***θ***_Γ_], e.g., 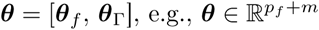.

In this work, we make several simplifying assumptions to the model in Eqs (1-2). First, we assume that the measurement function *h*(· ; ·) is linear and that all the parameters in ***θ***_*h*_ are known, so Eq (2) becomes

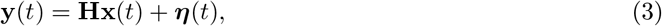

with **H** ∈ ℝ^*m×d*^. Second, we assume that the initial condition **x**(*t* = 0) = **x**_0_ is known, so it is excluded from parameter estimation.

Although Eqs (1-2) define a continuous-time dynamical system, we mostly deal with discrete observations. The set of *n* measurements 𝒴_*n*_ = {**y**_1_, …, **y**_*n*_} denotes the experimental data taken at time instances *t*_1_, ≤ *t*_2_, …, ≤ *t*_*n*_, where **y**_*k*_ = **Hx**_*k*_ + ***η***_*k*_. In this discrete setting, **x**_*k*_ is the state vector at time *t*_*k*_ and ***η***_*k*_ is an independent realization of the measurement noise. Additionally, the set of states at the discrete measurement times defined above is 𝒳_*n*_ = {**x**_1_, …, **x**_*n*_}. Note that the full (internal) states may not be available for estimation, only the measurements.

We are now ready to define the problem setting of the proposed parameter estimation framework. This work assumes that the model form is known and seeks to estimate the model parameters and associated uncertainties by learning a probability distribution for the parameters. The following problem statement formalizes the parameter estimation problem.

##### Problem 1.

*Given a known model form as in* *Eq* (1) *and a set of noisy, sparse and incomplete, experimental measurements* 𝒴_*n*_ = *{***y**_1_, …, **y**_*n*_*} at time instances t*_1_ *≤ t*_2_, …, *≤ t*_*n*_, *estimate the complete probability distribution, p*(***θ***|𝒴_*n*_), *for the model parameters* ***θ*** = [***θ***_*f*_, ***θ***_Γ_].

The next section outlines the structural identifiability and global sensitivity analyses performed to reduce the dimension of ***θ*** before estimating parameters.

### 2.3. Global structural identifiability analysis with the structural identifiability analyzer (SIAN)

Structural identifiability analysis determines if parameters can be uniquely estimated from the available measurement function [29]. Structural identifiability is a mathematical property of the model itself and does not consider the quality or quantity of the available experimental data. The following definition from [60] provides an intuitive condition for global structural identifiability and can be shown to be equivalent to alternative definitions [33, 36].

#### Definition 1

(Global structural identifiability [29, 61]). *A parameter θ*_*i*_ *is globally structurally identifiable if, for all times t >* 0,

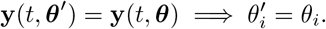

*Global* structural identifiability, as in Definition 1, implies that a parameter *θ*_*i*_ can be uniquely identified from data [36]. Alternatively, a parameter may be *locally* structurally identifiable if the condition in Definition 1 only holds in the local neighborhood, *v*(*θ*_*i*_), of parameter space around *θ*_*i*_, e.g., for *θ*_*i*_ ∈ *v*(*θ*_*i*_) [29, 33, 36]. We choose to use the differential algebra and power series-based approach given in [36, 52] to assess global structural identifiability. An overview of alternative approaches for structural identifiability analysis is provided in [29, 33, 36, 62] and the references within.

The SIAN (Structural Identifiability ANalyser) software [52] provides a numerical implementation of the approach proposed in [36]. The mathematical details of the approach are beyond the scope of this paper and out provides [36]; however, we provide a brief overview (see also, supplemental material of [52]) of the algorithm [36, 52]. The SIAN algorithm for assessing global structural identifiability uses a combination of symbolic and probabilistic computation. First, SIAN uses Taylor expansions of the model equations to obtain a polynomial representation of the system. Second, the algorithm truncates the polynomial system to produce a minimal system containing all parameter identifiability information. Third, SIAN solves the identifiability problem for a single parameter set that is randomly selected to guarantee correctness up to a user-specified probability level, *p* (see Theorem 5 in [36]). Fourth, the algorithm uses the results in the third step to separate the parameters into globally identifiable, locally identifiable, and nonidentifiable sets. SIAN is implemented in Maple (Maplesoft, Waterloo, ON) and Julia (The Julia Project [63]).

In this work, we use the Julia implementation of the SIAN algorithm with the default probability of correctness, *p* = 0.99 (available at https://github.com/alexeyovchinnikov/SIAN-Julia). However, the algorithm cannot guarantee this probability of correctness because we set the additional `p_mod` parameter to 2^29^ *−* 3 to enable the algorithm to run faster [64]. Any parameters that are not globally structurally identifiable are fixed to nominal values informed by the available literature following identifiability analysis. While these parameters may convey meaningful biological information, nonidentifiability implies that it is mathematically impossible to identify them from the available data. Next, global sensitivity analysis is used to further reduce the set of identifiable parameters.

### 2.4. Variance-based global sensitivity analysis

Parametric sensitivity analysis quantifies the contributions of the model parameters to variations in the model output [19, 37]. Specifically, global sensitivity analysis aims to quantify the effects of the model parameters on the output quantity of interest over the entire parameter space [19, 37] and is well suited for studying parameters in nonlinear models [37, 32]. This work applies Sobol’s method [53] for variance-based sensitivity analysis because it provides a quantitative output to rank parameters and leverages the prior distributions defined for the parameters. Sobol sensitivity analysis decomposes the variance of model outputs based on contributions from individual parameters and interactions between parameters [37, 53]. The total variance of the output quantity *f* (***θ***) is

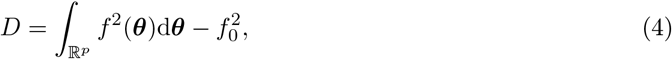

where 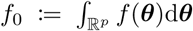 is the mean of the output. The following definition for the analysis of variance (ANOVA) representation provides an expansion for the output variance in a high-dimensional representation (HDMR), also known as a Sobol representation [19, 53].

#### Definition 2

(Analysis of variance (ANOVA) representation [19, 53, 37]). *The ANOVA expansion states that the output function f* (***θ***), *for* ***θ*** ∈ ℝ^*p*^ *defined as* ***θ*** = [*θ*_1_, *θ*_2_, …, *θ*_*p*_], *can be represented as*

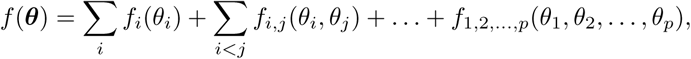

*where the zero-, first-, and second-order terms are defined recursively as*

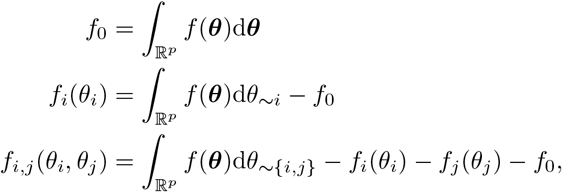

*respectively. The recursion is extended further for increasing numbers of parameters to compute higherorder terms. This definition assumes that the contribution terms are orthogonal (see Def 1 in [53]), and the notation ∼ i refers to the set excluding index i, for example:* d*θ*_*∼ i*_ = *{*d*θ*_1_, …, d*θ*_*i−* 1_, d*θ*_*i*+1_, …, d*θ*_*p*_*}*.

The ANOVA representation (see Def 2) expands the total variance, Eq (4), as

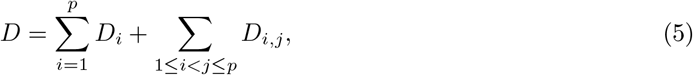

where the variances *D*_*i*_ and *D*_*i,j*_ are

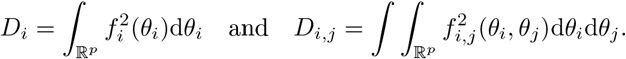

Note that it is possible to compute higher-order variances by increasing the dimension of the integral and following the recursion in Def 2, however we limit our discussion to second-order or lower variances for brevity. The first and second-order Sobol sensitivity indices are then defined using the variance terms in Eq (5) as

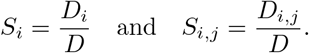

The first-order sensitivity index *S*_*i*_ quantifies the fraction of the total variance attributed to parameter *θ*_*i*_, and the second-order sensitivity index *S*_*i,j*_ quantifies this for the interactions between *θ*_*i*_ and *θ*_*j*_. Lastly, the *total-order sensitivity* is

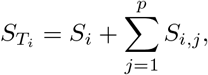

which quantifies all contributions from parameter *θ*_*i*_ on the output variance.

This work uses the UQLab toolbox [65, 66] to perform Sobol sensitivity analysis in Matlab (MathWorks, Natick, MA) and the `DifferentialEquations.jl` package [67] for analysis in Julia (The Julia Programming Language https://julialang.org) [63]. Both softwares use Monte Carlo estimators (see [19, 37] and references therein for details) to compute Sobol sensitivity indices from parameter samples. In Matlab, unless otherwise specified, first and total sensitivity indices are computed using Sobol pseudo-random sampling (e.g. `SOpts.Sobol.Sampling = ‘sobol’`) and the default estimator (`SOpts.Sobol.Sampling = ‘janon’`), see [66] for details. Additionally, the number of samples, `SOpts.Sobol.SampleSize`, is set specifically for each problem in section 3. In Julia, we perform similar Sobol sampling with `QuasiMonteCarlo.jl` (https://github.com/SciML/QuasiMonteCarlo.jl) with the `SobolSample()` sampler, and the default estimator for the sensitivity indices, e.g. `Ei_estimator` set to `:jansen1999`.

Sobol sensitivity analysis computes sensitivity indices for single scalar output quantities *x* rather than an entire trajectory. While it is possible to compute sensitivity indices at each time point in the trajectory and analyze the time series of indices, performing sensitivity analysis on specific quantities of interest is more interpretable. This work defines quantities of interest (QoI) for each problem that capture the biologically relevant information in a trajectory, such as the steady state value (at the final time in a trajectory). Parameters are ranked by the sensitivity indices for the given QoI. For simplicity, parameters are separated where there is a pronounced decline in the sensitivity index value. If this separation remains ambiguous, a threshold is chosen, for example, 0.1 ≤ *S*_*i*_, and any parameters whose sensitivity indices are above the threshold are considered for estimation. All parameters with *large* sensitivity indices are left free for estimation and the remaining parameters are fixed to nominal values. The next section recasts the parameter estimation problem, Problem 1, in the Bayesian framework to estimate the remaining free parameters.

### 2.5. Bayesian parameter estimation

Bayesian parameter estimation solves Problem 1 by considering the model parameters, ***θ***, as random variables that do not have a pre-specified parametric distribution. The approach characterizes the posterior probability distribution for the parameters *p*(***θ***|𝒴_*n*_) conditioned on a given dataset 𝒴_*n*_ and provides the *best guess* probability distribution for ***θ*** *given the data*. We can use Bayes’ rule to express the posterior distribution as

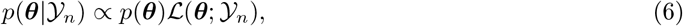

where *p*(***θ***) is known as the prior distribution, and ℒ (***θ***; 𝒴_*n*_) is the likelihood function (see Fig 1.B2 for a visual representation of Bayes’ rule).

Intuitively, Bayes’ rule updates our *best guess* about the distribution of the model parameters as new data is being incorporated. The prior distribution *p*(***θ***) represents the *best guess* before any data are collected and encodes any assumptions on the parameters. For instance, the prior may convey the physiological ranges for parameter values or may weigh known values more heavily (see section 2.8). The likelihood function ℒ (***θ***; 𝒴_*n*_) = *p*(𝒴_*n*_ |***θ***) updates our *belief state* by measuring the misfit between the data and model predictions of a specific parameter set. Parameter sets that are more likely to occur will produce model predictions that better match the data and thus have larger likelihood values. For example, although the prior in Fig 1.B2 places more probability on smaller values of *θ*, the likelihood in Fig 1.B2 places more probability mass towards larger values. It is important to note that evaluation of the likelihood function requires model simulations. For example, assuming Gaussian measurement noise with zero mean, a possible likelihood function is

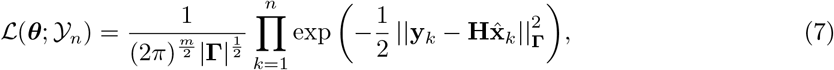

where 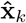 is the predicted state at time *t*_*k*_ with the parameters ***θ***, | *·* | denotes the matrix determinant, and the **C**-weighted norm is defined as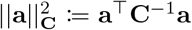. Lastly, the posterior distribution conveys the *best guess* after collecting and incorporating the data 𝒴_*n*_ into the statistical model and can be further refined if more data are included. In Fig 1.B2, the posterior illustrates how the likelihood function re-weights the prior to update our *belief state*.

A fundamental difficulty in Bayesian parameter estimation is that Bayes’ rule only enables evaluating the posterior distribution at specific points in parameter space. This is in contrast to, for example, the formula for a Gaussian distribution (with mean *µ* and standard deviation *σ*)

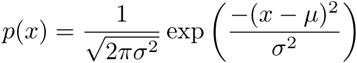

that can be analytically evaluated at all values of *x*. Therefore, parameter samples are drawn from the posterior help to characterize the distribution over the entire parameter space. Markov chain Monte Carlo (MCMC) algorithms enable sampling from arbitrary distributions, such as the posterior distribution (see section 2.9 for details). Before performing Bayesian estimation the next section introduces the constrained interval unscented Kalman filter Markov chain Monte Carlo (CIUKF-MCMC) algorithm that accounts for uncertainty in the model and the data.

### 2.6. Constrained interval unscented Kalman filter Markov chain Monte Carlo (CIUKF-MCMC)

A framework for complete uncertainty quantification of dynamical models in systems biology must account for uncertainty in the model form, parameters and noisy data. In [26], Galioto and Gorodetsky suggest adding a process noise term to Eq (1) to account for model form uncertainty in the system. Following this suggestion, the model in Eqs (1-2) is recast as a discrete time stochastic process

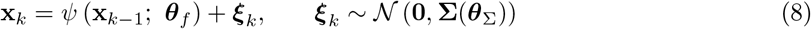

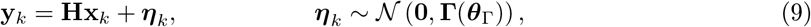

where *k* is the discrete time index for *t*_*k*_, and *ψ*(*·*; *·*) is the discrete state propagator that evolves the state from time *t*_*k−* 1_ to time *t*_*k*_. Additionally, ***ξ***_*k*_ and ***η***_*k*_ are Gaussian process and measurement noise (stochastic noise processes) with covariances **Σ**(***θ***_Σ_) and **Γ**(***θ***_Γ_), respectively. The discrete state propagator *ψ*(·; ·) in Eq (8) is the discrete operator that evolves the state according to the differential equation in Eq (1), and, for example, could be a forward-Euler finite-difference approximation. This work deploys the Matlab function `ode15s()` to construct a state propagator that guarantees the necessary stability to handle systems biology models.

Bayesian estimation of the model parameters, ***θ*** = [***θ***_*f*_, ***θ***_Σ_, ***θ***_Γ_], of the extended system in Eqs (8-9) accounts for uncertainty in the data, model, and parameters. The introduction of process noise increases the dimension of the parameters to estimate by requiring estimates for *θ*_Σ_. Further, the addition of stochastic process noise means the state variables are random variables; Bayes’ rule for this system becomes

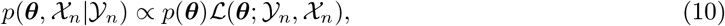

to account for the additional uncertainty in the states. The key step of the UKF-MCMC algorithm is the marginalization of uncertainty in the states out of Eq 10 to enable estimation of the parameter posterior distribution [26].

The UKF-MCMC algorithm begins by constructing an expression for the joint likelihood of the states and the parameters, ℒ (***θ***; 𝒴_*n*_, 𝒳_*n*_). Two probability distributions implied by the stochastic system in Eqs (8-9) are needed to define an expression for the joint likelihood. First, the probability of the current state **x**_*k*_ given the past state **x**_*k−* 1_ is

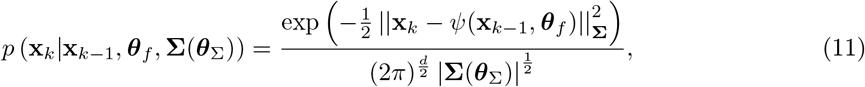

where the norm 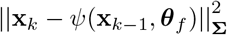 quantifies the misfit between the past state and the predicted current state. Next, the probability of a measurement **y**_*k*_ given **x**_*k*_ is

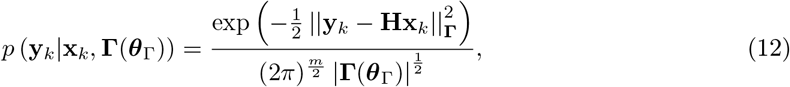

where the norm 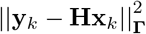 quantifies the residual between the measurement and the true states. By combining Eq (11) and Eq (12) the joint likelihood is

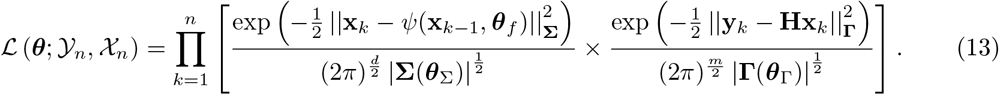

Marginalizing out the uncertain states by integration yields the likelihood for the uncertain parameters

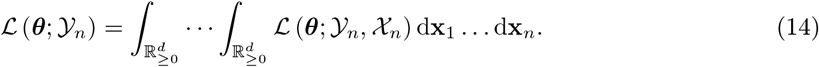

However, there is no obvious computationally tractable approach to integrate over a set of uncertain states directly. Theorem 1, stated below, provides a recursive algorithm to marginalize the states out of the likelihood, e.g., to perform the integration in Eq (14). Although Theorem 1 assumes that the initial condition is uncertain (and it is therefore estimated), we do not use that estimate in this work, as we start with a known initial condition **x**_0_.

#### Theorem 1

(Marginal likelihood (Theorem 1 of [26] and 12.1 of [68])). *Let 𝒴*_*k*_ *denote the set of all observations up to time k as defined in section 2.2. Let the initial condition be uncertain with distribution p* (**x**_0_|***θ***). *Then the marginal likelihood is defined recursively in three stages:*

*for k* = 1, 2, …

1. *Predict the new state from previous data*

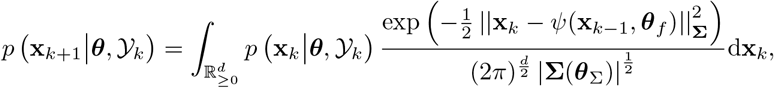
2. *update the prediction with the current data*

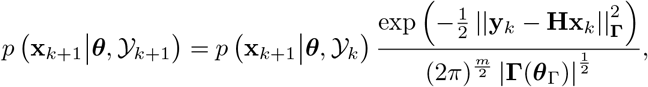
3. *and marginalize out uncertainty in the states*

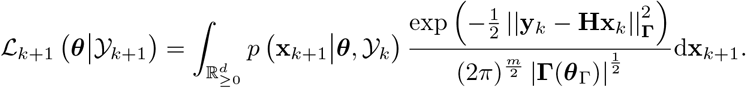

The recursion defined in Theorem 1 closely resembles a Bayesian filter [68]; thus, it is evaluated it with Kalman filtering algorithms [26]. For linear models, the standard linear Gaussian Kalman filter can be used to evaluate the recursion (see Algorithm 2 in [26]). However, exact solutions to the recursion are not possible if the model or measurement processes are nonlinear. Therefore, approximations such as extended Kalman filters, unscented Kalman filters, or ensemble Kalman filters [26, 68] must be employed. The original implementations of the UKF-MCMC algorithm use the UKF [69] for this approximation because the UKF is generally stable and can handle nonlinear models and measurement processes [26, 51]. However, the UKF is not suitable for systems biology models because it ignores constraints on the state variables, such as the nonnegativity of chemical concentrations. Section 2.7 describes the constrained interval unscented Kalman filter (CIUKF) [54] implemented in this work to enforce state constraints during filtering. Thus, we refer to the UKF-MCMC from [26] that uses the constrained interval unscented Kalman filter [54] as CIUKF-MCMC.

### 2.7. Constrained interval unscented Kalman filter (CIUKF)

We implement the constrained interval Unscented Kalman filter (CIUKF) [54, 70] algorithm to enforce all state constraints in CIUKF-MCMC. This algorithm assumes that the state is subject to an interval constraint **x**_LB_ ≤ **x** ≤ **x**_UB_. We only seek to enforce nonnegativity in systems biology, so the interval constraint is **0** ≤ **x** ≤ **∞**. We choose the CIUKF over alternative state constrained Kalman filters [54, 71] because it enforces constraints in both the predict and update steps of the algorithm and retains the same structure as the standard UKF [68]. We outline the steps of the CIUKF below.

The CIUKF algorithm predicts the state **x**_*k*+1_ from all preceding measurement data **y**_1_, …, **y**_*k*_. Following the structure of the linear Kalman filter, CIUKF is a recursive algorithm that iterates over all data and performs prediction and update steps at each time point [68]. For simplicity, we outline a single iteration of the CIUKF that moves the state forward in time from **x**_*k*_ to **x**_*k*+1_. Let 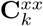 be the state covariance matrix at time *t*_*k*_, **Σ**(***θ***_Σ_) be the process noise covariance matrix, **Γ**(***θ***_Γ_) be the measurement noise covariance matrix, and ***θ***_*f*_ be the model parameters. There are two steps to the CIUKF, prediction, and update.

First, the prediction step uses the the interval constrained unscented transform [70, 54] (the state-constrained equivalent to the unscented transform [72, 69]) to predict the state and its covariance matrix at the next time point after propagation by the nonlinear model, e.g., Eq (8). The interval constrained unscented transform constructs a set of sigma points that capture the covariance 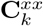 at time *t*_*k*_. Each sigma point is propagated in time by the nonlinear model to approximate the new state and its covariance at the next time *t*_*k*+1_. The set of 2*d* + 1 sigma points, 𝒳, is given by

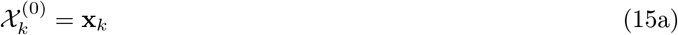

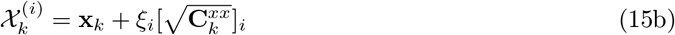

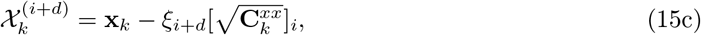

where *w*_*i*_ is the *i*th coefficient, [**A**]_*i*_ is the *i*th column of 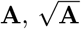 is the matrix square root of **A**, and *i* = 1, … *d*. The coefficients *ξ*_*i*_ control the distances of the sigma points around the initial state **x**_*k*_ and are chosen to ensure that no sigma points violate the state constraints. They are

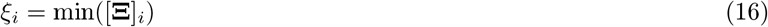

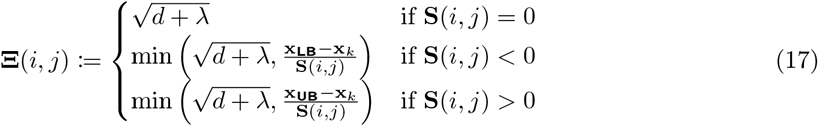

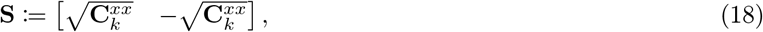

where *λ* is a parameter of the algorithm. Alternatively, in the standard unscented transform, the coefficients are all equal to 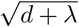 [69, 72]. Next, a set of weights, *w*_*i*_, are assigned to each sigma point as

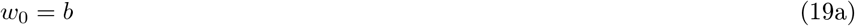

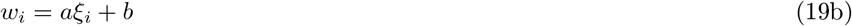

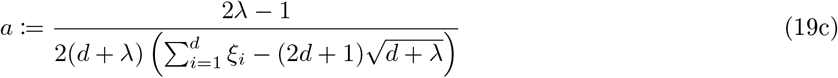

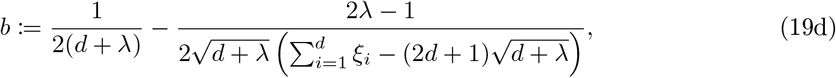

where *ξ*_*i*_ are as defined in Eqs (16-18). Importantly the sum of the weights equals one, 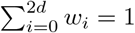. The prediction step then uses the nonlinear model, Eq (8), to propagate each sigma point forward in time

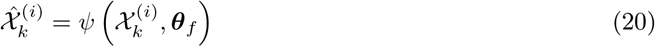

for *i* = 0, …, 2*d*. The prediction mean 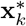 and covariance 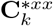 are then respectively computed as

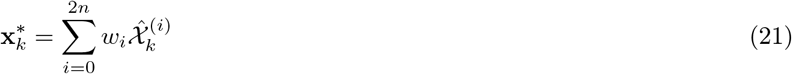

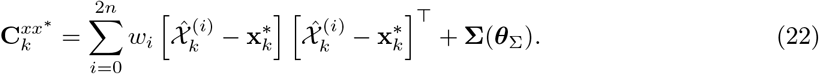

Equations (15)-(22) describe the constrained interval unscented transform that approximates the mean and covariance of the state after propagation by the nonlinear model. The prediction mean, Eq (21), and covariance, Eq (22), provide the predicted state and its covariance, respectively, that are then updated using the available data **y**_*k*_.

The update step begins by constructing a new set of sigma points centered around 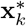, where

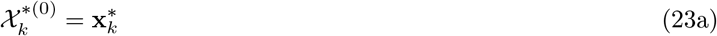

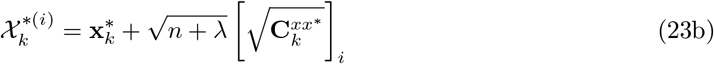

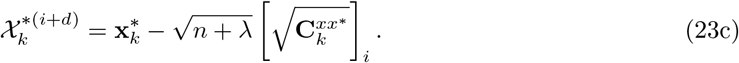

Additionally, a new set of weights are

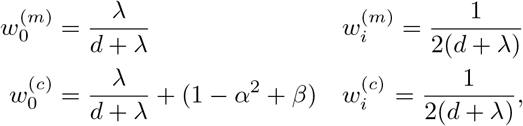

where 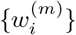 are used to compute the mean and 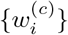 are used to compute the covariance matrix. These weights are indeed equal for all sigma points and are equivalent to those used in the standard UKF. Next, the measurement function is applied to each sigma point, yielding a set of predicted measurements

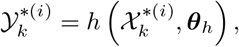

where the measurement function *h*(*·*; *·*) is possibly nonlinear with parameters ***θ***_*h*_. Then the mean and covariance matrices of the predicted measurements are computed with the weighted sums,

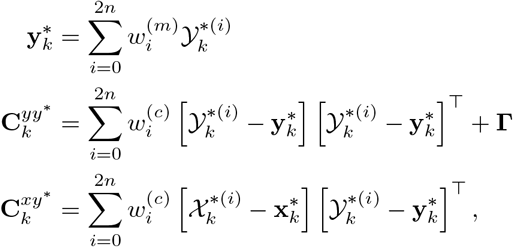

and the Kalman gain is

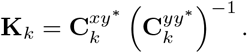

Lastly, the updated state **x**_*k*+1_ is found by solving the following constrained nonlinear optimization problem,

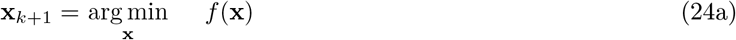

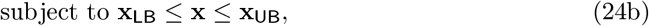

where the objective function is

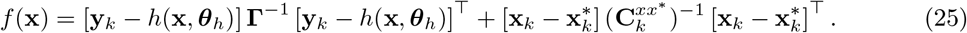

This optimization problem can be solved in Matlab using the `fmincon()` optimizer. Additionally, the updated covariance matrix is given by

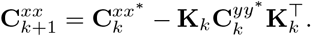

In offline state estimation problems, such as CIUKF-MCMC, this filter is iterated over all available data, e.g. from time *t*_0_ to time *t*_*n*_ if *n* data points are available [26, 51].

In practice, the CIUKF algorithm is substantially more compute-intensive than the standard UKF [54] because the CIUKF update step involves solving a constrained nonlinear optimization problem, e.g., Eq (24). However, the objective function in Eq (25) can be simplified given the linear measurement assumptions made in section 2.2. The simplified objective function becomes

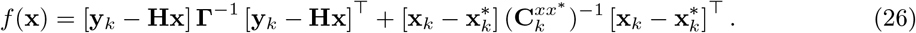

Expanding this and recognizing that minimizing *f* (*x*) = *y*(*x*) + *b* is equivalent to minimizing *f* (*x*) = *y*(*x*), gives

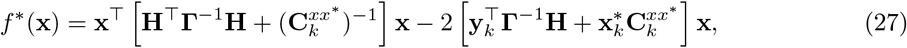

which is equivalent objective function to Eq (26) and is a quadratic form. Thus, the constrained optimization problem in Eq (24) becomes a quadratic program when using Eq (27) as the objective function. We use the `quadprog()` function in Matlab to solve the quadratic program with the `‘Algorithm’` option set to `‘trust-region-reflective’`. We keep all other settings for the solver as defaults. As expected, solving the quadratic program is substantially more efficient than solving the general nonlinear problem.

Throughout this work, we set *λ* = 1, *α* = 1 *×*10^*−* 3^, *β* = 1. Furthermore the Cholesky decomposition, **A** = **LL**^T^, is used to compute the matrix square roots in Eq (15) and Eq (23), because covariance matrices are always positive definite. Two additional numerical steps are taken to ensure all covariance matrices remain symmetric positive definite. First, *ϵ***I** is added to all computed covariance matrices, **P**^*∗*^ = **P** + *ϵ***I**, where *ϵ* = 1 *×*10^*−* 10^, and **P** is a generic covariance matrix. Second, computing 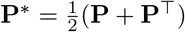 enforces symmetry of the covariance matrix. The next section discusses how to choose prior distributions for Bayesian estimation.

### 2.8. Prior distribution

The prior distribution, *p*(***θ***) encodes our *belief state* about parameters before collecting data and performing Bayesian estimation [19, 27]. The form of the prior distribution allows can be specified to incorporate varying levels of prior knowledge into our models. If the values of a parameter are known, informative priors can be used to shift to the possible values for that parameter towards the known values [27]; for example, a log-normal prior distribution can be used to center the prior around experimental measurements of a parameter [73]. However, the Bayesian inference literature often warns that informative priors should only be used in combination with good information on the parameter values [27] as it can take a large amount of data to overcome a “bad” prior. Alternatively, noninformative or weakly-informative priors reflect a lack of good prior knowledge about the parameters. For example, if we only know a parameter’s physiological ranges, we could construct a uniform prior that states there is an equal probability of any parameter value within this range. Noninformative or weakly-informative priors rely on the data to provide information on the parameters, so the choice of such priors is often safer when prior knowledge of the parameters is limited [27].

In applying the CIUKF-MCMC algorithm, this work constructs prior distributions for two sets of parameters, the biological model parameters, and the noise covariance parameters. We choose to use uniform priors for the biological model parameters, ***θ***_*f*_, to replicate the typical modeling setting where only the possible ranges for model parameters are known. Supplemental Tables in Appendix A list the upper and lower bounds of all biological model parameters. Furthermore, we follow the choices in [26] and use right-half-normal priors for the measurement and process noise covariance parameters, ***θ***_Σ_ and ***θ***_Γ_, respectively (this is further motivated in section 7.1 of [74]). The choice of covariance and upper bound of these priors was found to significantly affect the convergence of MCMC sampling. Thus, the prior distributions for the noise covariance parameters were scaled to match the respective state variable. For example, the prior for a measurement noise covariance 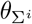 that corresponds to measurements 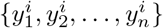 would be a right-half-normal distribution with mean zero and standard deviation equal to a fraction of the standard deviation 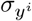 of the data. The prior is then 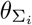 Right-Half-Normal 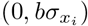 truncated to 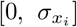, where *b <* 1 and 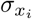 is the standard deviation of the state *x*_*i*_. We chose 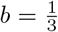 unless otherwise specified. These choices are motivated by the observation that measurement noise and process noise covariances will always be smaller than or equal to the covariance of the available data if there is any meaningful information in the data. The next section introduces MCMC sampling to characterize posterior distributions that use CIUKF to approximate the likelihood function.

### 2.9. Markov chain Monte Carlo sampling

Markov chain Monte Carlo (MCMC) algorithms enable sampling from arbitrary probability distributions [19, 75, 76, 77]. The key idea of MCMC is to construct a Markov chain of samples ***θ***^1^, ***θ***^2^, …, ***θ***^*N*^ whose distribution converges to the target distribution, *π*(***θ***), that we wish to sample [27, 76]. We apply MCMC sampling to Bayesian parameter estimation by constructing a Markov chain where the target distribution is the posterior distribution, that is *π*(***θ***) = *p* (***θ***|𝒴_*n*_). In this work, we use two MCMC sampling algorithms, delayed rejection adaptive Metropolis (DRAM) [58] and affine invariant ensemble sampler (AIES) [59]. These samplers build upon the classical Metropolis-Hastings algorithm [78, 79] that we introduce in section 2.9.1. We outline DRAM in section 2.9.2 and AIES section 2.9.3. Lastly, we discuss convergence assessment with the integrated autocorrelation time [59] in section 2.9.5. We focus these discussions on the practical aspects of MCMC and refer the reader to [19, 27, 76] for additional theoretical details.

#### 2.9.1. Metropolis-Hastings

The Metropolis-Hastings (MH) algorithm [78, 79] constructs a Markov chain whose probability distribution is guaranteed to converge to the target distribution and forms the foundation for a large family of MCMC samplers [27, 76]. The MH algorithm consists of two steps, a proposal and an accept-reject step, repeated to draw the set of samples ***θ***^1^, ***θ***^2^, …, ***θ***^*N*^. The Markov chain starts with an initial sample ***θ***^0^ that the user chooses. We outline one iteration of the MH algorithm moving from step *i* to step *i* + 1.

The first step, called the proposal step, in Metropolis-Hastings is to propose a new sample ***θ***^*∗*^. We draw ***θ***^*∗*^ from the proposal distribution *q*(***θ***^*∗*^ |***θ***^*i*^). The proposal distribution is specific to each MCMC algorithm, but, for example, could be a normal distribution centered around the previous sample such that ***θ***^*∗*^ *∼* 𝒩 (***θ***^*i*^, *σ*^2^**I**) where *σ* is specified (this is random walk Metropolis (RWM) [76]).

Next, the accept-reject step decides if the proposal is accepted, set ***θ***^*i*+1^ = ***θ***^*∗*^, or rejected, set ***θ***^*i*+1^ = ***θ***^*i*^. In the MH algorithm, this decision is probabilistic, e.g., the proposal is accepted or rejected with probability *α*(***θ***^*∗*^|***θ***^*i*^). The acceptance probability is

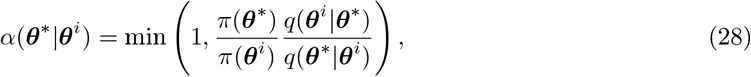

which guarantees that the stationary distribution of the samples (the distribution that the samples converge to in the infinite sample limit) equals the target distribution [27, 76]. These steps, propose and accept-reject, are repeated until the distribution of the set of samples has *converged* to its stationary distribution. Although convergence is guaranteed in the infinite sample limit [27, 76], assessing convergence in practice is nontrivial. We chose to use an approach from [59] that uses the integrated autocorrelation time to asses convergence as outlined in section 2.9.5 (see [19, 27, 76] for additional approaches).

#### 2.9.2. Delayed rejection adaptive Metropolis (DRAM)

The delayed rejection adaptive Metropolis (DRAM) algorithm is based on the random walk Metropolis algorithm and combines delayed rejection with adaptive Metropolis [58]. DRAM closely follows the Metropolis-Hastings algorithm but specifically uses a Gaussian proposal distribution such that ***θ***^*∗*^ ∼ 𝒩(***θ***^*i*^, **C**), where **C** is the covariance matrix. Delayed rejection (DR) [58, 80] and adaptive Metropolis (AM) [58, 81] are two modifications to the random walk Metropolis algorithm that help to improve the convergence rate of the Markov chain.

First, delayed rejection adds an additional round of Metropolis-Hastings (propose and accept or reject) if the first proposal is rejected. That is, if ***θ***^*∗*^ is rejected a new proposal, ***θ***^*∗*2^ *∼ q*_2_(***θ***^*∗*2^|***θ***^*∗*^, ***θ***^*i*^) is drawn, where *q*_2_(*·*) is the new proposal distribution. The new proposal distribution is also Gaussian, how (ever the covariance is scaled to a fraction of the original proposal covariance, e.g. *q*_2_(***θ***^*∗*2^|***θ***^*∗*^, ***θ***^*i*^) = 𝒩 (***θ***^***i***^, *γ***C**), where *γ <* 1 and is a tuning parameter of the algorithm [58]. Thus the second proposal is closer to the previous point and is more likely to be accepted. DRAM evaluates the new proposal with a Metropolis-Hastings accept-reject step with the acceptance probability

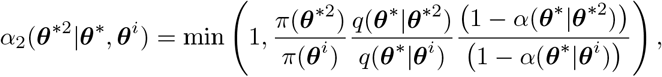

where *α*(· | · ′) is as defined in Eq (28). Although most implementations impose a single DR step [19], delayed rejection can be repeated more than once, where the proposal distribution and acceptance probability are modified accordingly at additional each round.

Adaptive Metropolis acts separately from delayed rejection and aims to move the proposal distribution closer to the target distribution [58], by replacing the covariance matrix of the proposal, **C**, with the covariance matrix of the samples. In practice, adaptation begins after a set number of *i*_0_ samples have been drawn and updates the covariance matrix at every step as

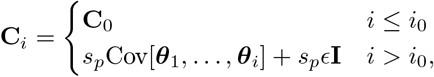

where **C**_0_ is the initial covariance matrix, *s*_*p*_ is an algorithm tuning parameter that is often set to *s*_*p*_ = 2.38^2^*/p* [58, 75] where *p* is the dimension of ***θ***, and *ϵ* is a small positive number.

We implement the DRAM algorithm with a single round of delayed rejection following [26] with *γ* = 0.01, *i*_0_ = 200, and *ϵ* = 1 *×*10^*−* 10^. Furthermore, the initial covariance matrix **C**_0_ is tuned to accept between 20 - 40% of proposals. We refer the reader to [19, 58] for further details on DRAM and tuning the initial covariance matrix. Additionally, each Markov chain is initialized to the MAP point by using `fmincon()` (default options except: `‘UseParallel’` set to `true` and `‘MaxFunctionEvaluations’` set to 10, 000) in Matlab to minimize the negative log-posterior (equivalent to maximizing the posterior) as in [26].

#### 2.9.3. Affine invariant ensemble sampler (AIES)

While random walk Metropolis-based algorithms such as DRAM can adequately sample complex posterior distributions, these methods will show very slow convergence when the target distribution is highly anisotropic [59]. The posterior distribution with CIUKF-MCMC in systems biology are anisotropic because the scales of model parameters can vary by several orders of magnitude, and the noise covariance parameters often have different scaling than the model parameters. Fortunately, AIES provides an algorithm to sample such anisotropic distributions [59] effectively. The motivation for affine invariance is that anisotropic distributions can be transformed to isotropic distributions with an affine transformation. Thus an algorithm that is invariant to such transformations will effectively sample an isotropic distribution when sampling an anisotropic distribution [59].

The AIES algorithm differs from DRAM and random walk Metropolis because it leverages an ensemble of Markov chains rather than a single chain. Each chain in the ensemble of *N*_*e*_ Markov chains is called a walker, and we denote the set of walkers at step *i* with 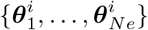 where the subscript is the walker index and the superscript is the step-index. Note that the number of walkers must be larger than the dimension of ***θ*** that is *N*_*e*_ *> p*, for ***θ*** ∈ ℝ^*p*^. At the end of *N* steps, a set of *N*_*e*_ Markov chains with *N* samples is obtained. The first chain in the ensemble is, for example,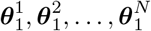. Note that the total ensemble will have *N ·N*_*e*_ samples. Each step of the AIES algorithm involves a proposal and a Metropolis-Hastings update for each chain in the ensemble. One iteration of these steps to move from *i* to *i* + 1 for a single walker, 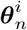, is outlined below and these steps are repeated to update the entire ensemble.

First, a proposal for the current walker 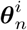 is chosen using the stretch move [59] that ensures affine invariance of the sampler. The stretch move proposes a new point that lies along the line

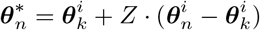

that connects the current walker 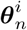 and another walker in 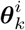 randomly chosen from the ensemble. Here, *Z* is a realization of *z ∼ g*(*z*), where

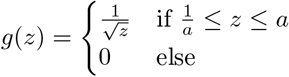

and *a >* 1 is an algorithm tuning parameter that the user must specify.

Second, the proposal is accepted or rejected using a Metropolis-Hastings-like accept-reject step. The acceptance probability is

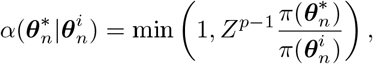

where *Z* is as defined above, and *p* is the dimension of ***θ***. This formulation of the acceptance probability guarantees convergence to the target distribution [59].

We use the Matlab implementation of the AIES algorithm [82] in the UQLab toolbox [65]. Unless otherwise specified, the ensemble size is *N*_*e*_ = 150 because we observed improved sampling with a large ensemble. To accelerate sampling, the likelihood is evaluated for each ensemble member in parallel using a parallel for loop (e.g., `parfor` in Matlab) with at most 24 parallel threads. Additionally, default value of *a* = 2 is used for the stretch move tuning parameter. Lastly, each Markov chain in the ensemble is initialized to a random point drawn uniformly over the support of the prior as is the default in UQLab [82].

#### 2.9.4. Markov chain burn-in

In MCMC, Markov chains (or ensembles of chains) often display an initial transient, called *burnin*, before converging to their stationary distributions [19, 27, 77]. Importantly, these samples during burn-in are not distributed according to the stationary distribution and should therefore be excluded from the final set of samples. Common practice in the MCMC literature is to simply discard these initial samples to remove the effects of burn-in [19, 27]. The choice of the burn-in length is often nontrivial and is best informed by an analysis of the Markov chain [77]. Unless otherwise specified the integrated autocorrelation time (described below) dictates the number of samples to discard as burn-in. Specifically, we compute the integrated autocorrelation time after collecting many samples and set the burn-in length to 5 - 10 times the computed value.

#### 2.9.5. Convergence assessment with the integrated autocorrelation time

A key challenge in Markov chain Monte Carlo sampling is determining the appropriate number of samples *N* to collect. In this work, we use the integrated autocorrelation time [59, 77] to determine when the Markov chain has approximately converged to its stationary distribution. We outline the theory and motivation behind the integrated autocorrelation time for a single Markov chain and refer the reader to [59] for a discussion of ensemble methods. The use of the integrated autocorrelation time is motivated by the typical use of MCMC sampling to compute an expectation

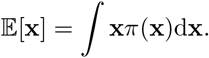

Given a Markov chain of length *N*, we can estimate the expectation with the Monte Carlo estimator

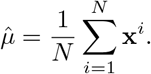

In general it is common to consider the variance of 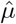, var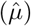, as the estimation error [59]. This variance is given by

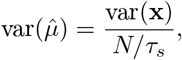

where the integrated autocorrelation time *τ*_*s*_ is given by

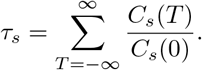

Here, the autocovariance function *C*_*s*_(*T*) with lag *T ∈* ℕ is given by

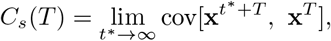

where *t*^*∗*^ ∈ ℕ. Thus, the Monte Carlo estimation error is proportional to integrated autocorrelation time for a fixed chain length. The integrated autocorrelation time can be interpreted as the time it takes for the samples in a Markov chain to become uncorrelated [77]. Additionally, the effective number of samples, *N*_eff_, can be defined using the integrated autocorrelation time as, *N*_eff_ = *N/τ*_*s*_.

In this work, we use the integrated autocorrelation time for two purposes. First, the computation of the integrated autocorrelation time is used to choose the correct burn-in length. We compute the integrated autocorrelation time after collecting many samples and then discard between 5 *−* 10 times *τ*_*s*_ as burn-in. Second, after discarding the initial samples, the integrated autocorrelation time helps to determine if enough samples have been collected, e.g., *N*_eff_ is large. Should the effective sample size be small, the MCMC sampler is run longer to collect more samples. We compute the integrated autocorrelation time using a Matlab function associated with [83] (available at https://www.physik.hu-berlin.de/de/com/UWerr_fft.m) with default algorithm parameters. We compute *τ*_*s*_ for each parameter and take the maximum of these values. For an ensemble, we compute the mean *τ*_*s*_ for the walkers of each parameter and take the maximum value of the means. After completing MCMC and evaluating the chains, the framework leverages the posterior samples to quantify uncertainty in model predictions.

#### 2.10. Ensemble simulation and output uncertainty analysis

Markov chain Monte Carlo sampling provides a set of samples, 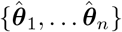, that converge in distribution to the posterior distribution (e.g. Fig 1.B3). One is often interested in how uncertainty in the parameter estimates, which is conveyed by the posterior distribution, propagates to uncertainty in the model predictions. Fortunately, an ensemble of simulations (see Fig 1.B1) distributed according to the posterior can be run using the posterior samples. This approach to uncertainty propagation is known as sampling-based uncertainty propagation [19] and is feasible because the simulation of dynamical systems biology models is computationally efficient. We refer the reader to [40, 84] for examples of sampling-based uncertainty propagation in systems biology.

Each simulation is run by solving the differential equation with the `ode15s()` integrator in Matlab with a unique sample from 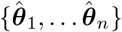 for the parameters. We use default tolerances for the integrator and supply the Jacobian matrix (specify the `‘Jacobian’` option) to improve computations. We compute the Jacobian by hand and evaluate it with the same parameter as the differential equation.

Following ensemble simulation, statistical analyses of the ensemble of predicted trajectories and any relevant quantities of interest (QoI) can be performed. For example, Fig 1.B2 highlights the uncertainty in the trajectory with 95% credible intervals to show the region where 95% of the trajectories fall. Additionally, we compute the statistics of relevant QoIs, such as the steady state values or limit cycle period, from the ensemble. The next section discusses how to choose single point estimates that best represents the estimated parameters.

### 2.11. Parameter point estimates from posterior samples

One often wants to compute a point estimate for each of the parameters in addition to characterizing the entire posterior distribution (black line in Fig 2). Common choices for point estimates in Bayesian statistics include mean, median, or mode of the posterior distribution [27]. Fig 2 highlights the mean and mode (denoted MAP), along with a secondary mode that may confound choosing a point estimator. The mode of the posterior distribution is a strong choice because it provides the most probable set of parameters and is often called the maximum *a posteriori* (MAP) point. However, the mean or median may provide better point estimates when the posterior is multimodal (there are multiple modes).

These point estimates are computed from posterior samples acquired via MCMC sampling. While sample statistics, such as the mean and median, are computed directly from the samples, computing the MAP point requires estimating the posterior of the probability density function and finding the maximal point. Direct computation of the sample mode will not yield the MAP point because the parameters are continuous random variables, so no two-parameter samples are expected to be identical. One approach to estimate the MAP point involves computing a histogram of samples and using the center of the bin with the most associated probability as the MAP. However, we found that the histogram of posterior samples is often noisy and can be sensitive to the bin size, so that the MAP estimate may be erroneous. We chose to take an alternative approach that fits a kernel density estimator [85] to approximate the posterior distribution and subsequently compute that MAP point. The kernel density estimator provides a non-parametric approximation of the posterior distribution and, intuitively, smooths the posterior histogram. We use the `ksdensity()` function for kernel density estimation in Matlab. The `‘support’` option is set to be the region bounded by the prior distribution and the `‘BoundaryCorrection’` option is set to use the `‘reflection’` method to account for these bounds. Lastly, we use the default values for the `‘Bandwidth’` unless otherwise specified. All other options are kept to the defaults as defined in the Matlab documentation. The MAP point is the point with maximum probability in the `ksdensity()` output. The next section moves from the details of CIUKF-MCMC and Bayesian estimation to outline how synthetic data is generated for the examples in this paper.

### 2.12. Synthetic data generation for numerical experiments

The synthetic data in this work aims to replicate noisy data found in biological experiments. We generate noisy synthetic data by drawing samples from deterministic model simulations and simulate measurement noise by adding independently and identically distributed (iid) perturbations to each sample. First, a nominal set of biological model parameters and an initial condition are chosen for data generation. These values become the *ground truth* for the estimation problem, and are informed by the available literature when possible. Next, a numerical solution to the system provides *true* trajectories of the state variables. Unless otherwise specified the `ode15s()` integrator in Matlab is used with default tolerances, and the Jacobian matrix is supplied (specify the `‘Jacobian’` option with the analytical Jacobian Matrix) to improve computations. We then apply the measurement function and sub-sample the *true* trajectory to simulate sparse sampling. Lastly, we corrupt the data by adding a realization of an iid noise stochastic process to each sample. To meet the assumptions of the CIUKF-MCMC algorithm (see Section 2.6) we use mean-zero, normally distributed perturbations with the diagonal covariance matrix, **Γ**. We chose the entries of **Γ** to be proportional to the variances of the respective state variables, e.g.

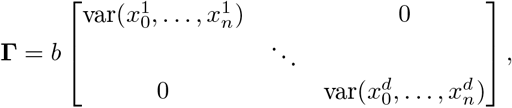

where *b* is a positive constant typically chosen to be less than one that controls the noise level. The last section discusses how to compute the amplitude and period of a limit cycle oscillation.

### 2.13. Limit cycle analysis

Limit cycle amplitude and period are used to characterize limit cycle oscillations for global sensitivity and output uncertainty analysis. These quantities are relevant in intracellular signaling because the strength and timing of signals, amplitude and period, respectively, are thought to encode different inputs [86]. The limit cycle amplitude, *y*_lca_, quantifies the difference between the maximum and minimum values of the oscillations and is defined as

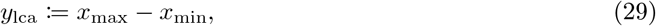

where *x*_min_ and *x*_max_ denote the minimum and maximum values of the state *x* over a single complete oscillation. Further, the limit cycle period, *y*_period_, is the time to complete an oscillation and is defined as

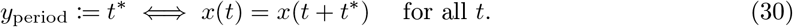

To compute these quantities we find trajectories that show limit cycle oscillations and then extract the two quantities of interest. This approach leverages the `findpeaks()` function that returns the locations of the local maxima (peaks) in a trajectory for these computations in Matlab. The `findpeaks()` can also find the local minima by applying it to the negative of the trajectory.

The first task in computing the limit cycle features is to detect actual limit cycle oscillations. A trajectory is discarded if it reaches a fixed point (steady state) when no peaks are detected (`findpeaks()` returns an empty set). Next, the difference between the heights of the identified peaks is used to discard trajectories that show decaying oscillations to a fixed point. A threshold on this difference, called for the `peakThreshold`, is set to 17.0 unless otherwise specified; a trajectory is discarded if its difference in peak values exceeds this threshold. Any remaining trajectories will show limit cycles or will contain numerical artifacts which are falsely detected as limit cycles.

Lastly, the limit cycle amplitude and period are computed. The limit cycle amplitude is the mean difference between each pair of detected peaks and minima that correspond to one oscillation of the limit cycle. The limit cycle period is then computed as the mean time between two peaks and the time between two minima. Trajectories with numerical artifacts are eliminated by discarding those with a limit cycle amplitude that is smaller than the `LCAminThresh`, with a range of limit cycle values greater than the `Decaythresh` or those that return no limit cycle period (e.g., the empty matrix on Matlab). Unless otherwise specified, we set the `LCAminThresh` to 1.0 and the `Decaythresh` to 5.0. These computations assume that the period of the limit cycle is stable and that no frequency modulation occurs.

## 3. Results

We applied the Bayesian parameter estimation framework to three different models that represent signal transduction cascades of increasing biological and mathematical complexity. Section 3.1 uses the first model, a simple kinetic scheme, to describe a series of computational experiments that illustrate the effects of measurement noise and data sparsity on estimation uncertainty (Fig 3). Next, we tested our framework on two models that are more representative of the nonlinearities and over-parameterization observed in systems biology models. Section 3.2 analyzes a representative model of the mitogen-activated protein kinase (MAPK) pathway [56] that exhibits multistability depending on the choice of model parameters. Finally, section 3.3 analyzes a simplified model of synaptic plasticity [57] that illustrates the effects of higher parameter uncertainty even in phenomenological biological representations.

**Figure 3:**
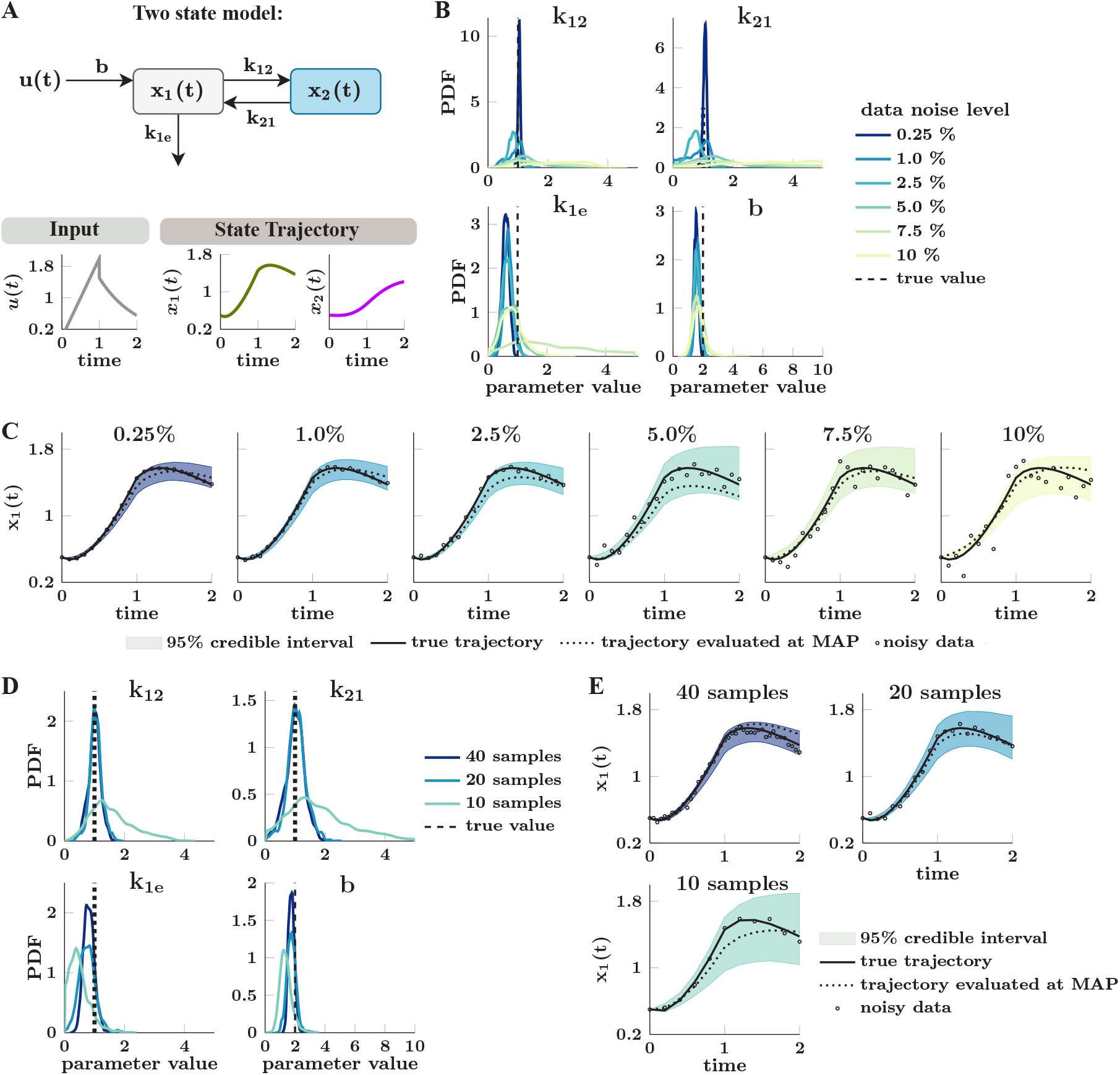
Parameter estimation for a simple two-state model. (**A**) *Top row:* Network diagram of the two-state model with states, *x*_1_(*t*), *x*_2_(*t*), input function *u*(*t*), and four unknown parameters, ***θ***_*f*_ = [*k*_1*e*_, *k*_12_, *k*_21_, *b*]. *Bottom row:* Trajectories of the input function *u*(*t*) and corresponding state trajectories. The input has at least one non-zero derivative to ensure that all model parameters are globally structurally identifiable following [55]. (**B**) Marginal posterior distributions of the model parameters show increasing uncertainty in the parameter estimates (e.g. widening and flattening) with increasing levels of additive normally distributed measurement noise with mean zero. We control the noise level by setting the noise covariances to the specified percentage of the standard deviation of each state variable. The dashed black vertical lines indicate each parameter’s nominal (true) value. Marginal posteriors are visualized by fitting a kernel density estimator to 20, 000 MCMC samples obtained using CIUKF-MCMC with the delayed rejection adaptive Metropolis (DRAM) MCMC algorithm after discarding the first 10, 000 samples as burn-in. (**C**) Posterior distributions of the trajectory of *x*_1_(*t*) reflect increasing parameter estimation uncertainty in panel B. The true trajectory (solid black line) shows the dynamics with the nominal parameters, dashed black lines show that trajectory with the most probable set of parameters (MAP point), and the empty circles show the noisy data at the specified noise level. The 95% credible interval shows the region between the 2.5th and 97.5th percentiles that contains 95% of the 5, 000 trajectories. (**D**) Marginal posterior distributions of the model parameters show increasing uncertainty (widening and flattening) with increasing data sparsity (fewer samples). We simulate data sparsity by sampling the simulation from 0 ≤ *t* ≤ 2 with three time steps, Δ*t* = 0.05 (40 experimental samples), Δ*t* = 0.1 (20 experimental samples) and Δ*t* = 0.2 (10 experimental samples). Marginal posteriors are fit to 20, 000 MCMC samples obtained as in panel B. (**E**) Posterior distributions of the trajectory of *x*_1_(*t*) reflect increasing parameter estimation uncertainty seen in panel (**D**).

### 3.1. Measurement noise and sparsity increase estimation uncertainty in a simple model of signal transduction

The first model we consider is a relatively simple two-state model from [55] shown in Fig 3.A. The governing equations for this model are

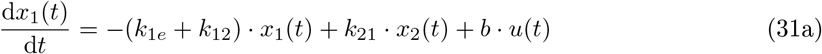

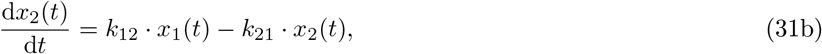

where the states are **x**(*t*) = [*x*_1_(*t*), *x*_2_(*t*)], the biological model parameters are ***θ***_*f*_ = [*k*_1*e*_, *k*_12_, *k*_21_, *b*], and *u*(*t*) is the input function. The input function (illustrated in Fig 3.A)

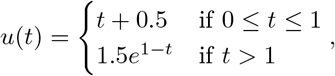

ensures that all four biological model parameters are structurally identifiable because the input function has at least one nonzero derivative [55]. Supplemental Table A.1 lists the nominal parameter values and the initial condition was **x**_0_ = [0.5, 0.5]. Sensitivity analysis was not performed on this model because the state variables are linearly dependent on the parameters.

Synthetic data with full state measurements (see section 2.12) was used to perform two parameter estimation experiments—one with increasingly noisy data and another with increasingly sparse data— to investigate how measurement noise and data sparsity affect parameter and output uncertainty. The noise levels of each dataset were controlled by taking a fraction of the maximum values of the true trajectory for the corresponding state variables. In each estimation experiment, we chose the prior distributions as outlined in section 2.8 and used CIUKF-MCMC (section 2.6) with DRAM (section 2.9.2) to draw 30, 000 posterior samples conditioned on this noisy data. To ensure sample counts were constant across noise and sparsity levels, we chose a constant burn-in length of 10, 000 samples to discard from every Markov chain (see section 2.9.4 for details). Supplemental Fig B.10 shows the Markov chains for the measurement noise experiments, and Supplemental Fig B.11 shows those for data sparsity experiments. The following remark highlights an important distinction between data points and MCMC samples.

#### Remark 1.

*Data points are different than MCMC samples. The experiments in this work produce at most 40 (simulated) data points, i.e*., *noisy measurements of the states, for parameter estimation. However, MCMC algorithms draw 10,000s-1,000,000s of sample parameter sets to characterize the posterior distribution, which requires evaluating the likelihood and, therefore, simulating the model*.

A first hypothesis we tested was that noisy data increases uncertainty because measurement noise limits the ability to constrain the dynamics of the state variables. To test this hypothesis, we performed Bayesian parameter estimation from synthetic data with increasing measurement noise (circle marks in Fig 3.C). We observed that the marginal posterior distributions in Fig 3.B (kernel density estimator fit to the posterior samples as in section 2.11) widen and flatten with increasing noise levels, indicating increased uncertainty. However, the most probable value for each parameter, the MAP point, lies close to the nominal parameter values (dashed lines in Fig 3.B) for every noise level, suggesting that the data provide information about the parameter irrespective of the noise level. Additionally, Fig 3.C shows that the width of the 95% credible interval for the dynamics of *x*_1_(*t*) grew as the noise increased from the lowest level (2.5%) to 5.0% and then remained similarly wide at the highest values (see Supplemental Fig B.9.A for the respective trajectories of *x*_2_(*t*)). While the uncertainty bound did not widen above the 5.0% noise level, the shape of the trajectories began to shift further from the truth (dotted line in Fig 3.C), indicating the estimates began to take on a bias. These experiments validated our hypothesis that even in a simple dynamical system measurement noise increases estimation uncertainty of kinetic parameters.

Next, we hypothesized that data sparsity (fewer data points) would increase the uncertainty in parameter estimates. To test this, we fixed the measurement noise to the 2.5% level (see Fig 3) and varied the number of measurements (e.g., the sampling rate) that were included in the data used for estimation. We tested three sparsity levels with 40 experimental samples (Δ*t* = 0.05), 20 experimental samples (Δ*t* = 0.1) and 10 experimental samples (Δ*t* = 0.2) over the simulation time 0≤ *t* ≤ 2. Fig 3.D highlights the widening of the estimated marginal posterior distributions for each model parameter as we decreased the number of data points (increased sparsity). Additionally, Fig 3.E shows that the increased parameter estimation uncertainty translates to increased uncertainty (wider 95% credible interval) in the trajectory of *x*_1_(*t*) (see Supplemental Fig B.9.B for the trajectories of *x*_2_(*t*)). In both of these experiments, the proposed uncertainty quantification framework qualitatively and quantitatively confirmed that increasing the noise or sparsity level increases estimation uncertainty.

### 3.2. Parameter estimation for a model of the MAPK cascade

We chose a simplified model of the highly conserved mitogen-activated protein kinase (MAPK; also known as the MEK/ERK cascade) signaling pathway [87] as a second test case for our parameter estimation framework. This pathway is known to exhibit bifurcations in its dynamical behavior [86, 88]; the system can reach a stable steady state or exhibit limit cycle oscillations. We focused on a phenomenological model of the MAPK pathway from [56] (see diagram in Fig 4.A) that includes the mixed feedback (negative and positive feedback) necessary to predict the range of dynamical behavior observed in experiments. This model has three states, *x*_1_(*t*), *x*_2_(*t*), *x*_3_(*t*) that correspond to phosphorylated RAF, MEK, and MAPK/ERK, respectively [89, 90] and 14 model parameters. The differential equations are

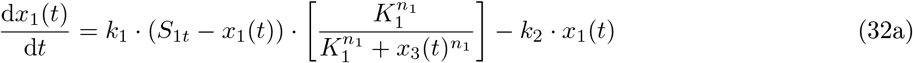

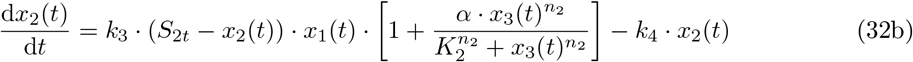

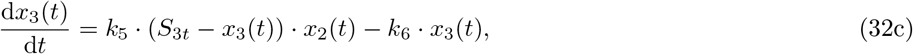

with the biological model parameters ***θ***_*f*_ = [*k*_1_, *k*_2_, *k*_3_, *k*_4_, *k*_5_, *k*_6_, *K*_1_, *K*_2_ *S*_1*t*_, *S*_2*t*_, *S*_3*t*_, *α, n*_1_, *n*_2_]. A previous analysis of the model in [56] found that it can predict three regimes of dynamical behavior that depend on the model parameters; these regimes are limit cycle oscillations, bistability and mixed multistability. Here we focus on using CIUKF-MCMC to estimate model parameters that produce two of the three dynamical regimes – bistability and limit cycle oscillations (see Fig 4.B for example trajectories). Supplemental Tables A.2 and A.3 list the nominal parameter values and initial conditions (as defined in [56],) respectively, used to produce each of these dynamics.

**Figure 4:**
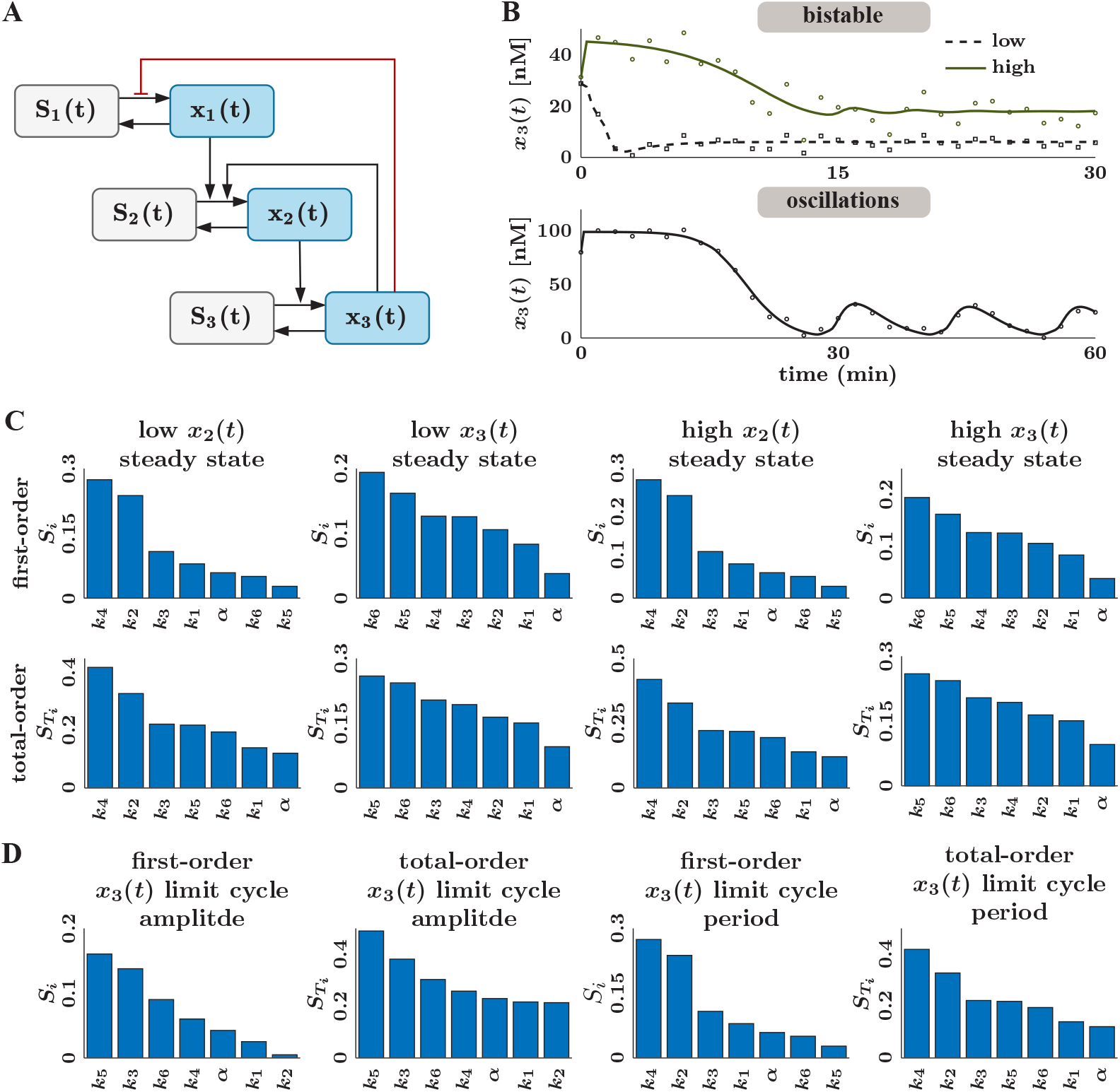
Parameter estimation for a simplified MAPK cascade that exhibits multistability. (**A**) Network diagram of the model of the core MAPK signaling cascade. The red line indicates inhibition; the black lines indicate activation. (**B**) Trajectories of *x*_3_(*t*) with the sets of nominal parameters that produce bistability (top) and limit cycle oscillations (bottom). The two dynamical regimes correspond to two different sets of nominal parameter values. The low (black dashed line) and high (solid green line) steady states are reached by manipulating the initial condition **x**_0_. The initial condition for the high steady state is **x**_0,high_ = [0.1245, 2.4870, 31.2623] and that for the low steady state is **x**_0,low_ = [0.0015, 3.6678, 28.7307]. (**C** and **D**) Sobol sensitivity for the MAPK model parameters. All parameters except the total concentrations, *S*_1*t*_, *S*_2*t*_ and *S*_3*t*_, and exponents, *n*_1_ and *n*_2_, are varied uniformly over the identified ranges (see Supplemental Table A.2) and 5000 samples are used for each parameter. (**C**) Sensitivity indices for bistable behavior dynamics. We use the steady state value of *x*_2_(*t*) and *x*_3_(*t*) for both the high and low steady states as quantities of interest. By selecting the two most sensitive parameters for the four quantities of interest we reduce the set of free parameters to ***θ***_*f*_ = [*k*_2_, *k*_4_, *k*_5_, *k*_6_]. (**D**) Sobol sensitivity indices for a set of free parameters that contribute to limit cycle behavior. We show the first-order sensitivity indices *S*_*i*_ and the total-order indices 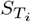 for the limit cycle amplitude and period of *x*_3_(*t*). We reduce the number of free parameters by selecting those with a first-order index greater than 0.1, *S*_*i*_ *>* 0.1, for the limit cycle amplitude or the period, e.g., ***θ***_*f*_ = [*k*_2_, *k*_3_, *k*_4_, *k*_5_].

First, we performed identifiability and sensitivity analysis to find the subsets of relevant parameters to estimate each dynamical regime. The SIAN software [52] (see Section 2.3) showed that 12 of the 14 biological model parameters are structurally identifiable from measurements of all three state variables; however, *K*_1_ and *K*_2_ are only locally structurally identifiable. SIAN cannot assess parameters that appear in an exponent [36, 52], so we fixed *n*_1_ and *n*_2_ to their nominal values listed in Supplemental Table A.2. To avoid global versus local identifiability complications, we fixed *K*_1_ and *K*_2_ to their nominal values. Additionally, we omitted the total concentration parameters, *S*_1*t*_, *S*_2*t*_, and *S*_3*t*_, from further analysis because we assume they would be specified according to the cell type that corresponds to the available data. As a result, we narrowed the free parameters down to a set of 9 parameters, from a set of 14 originally.

Next, we used global sensitivity analysis as described in Section 2.4 to further reduce the number of free parameters. We choose the quantities of interest for sensitivity analysis for the bistable and oscillatory regimes separately. The quantities of interest for the bistable regime are the steady state values of *x*_2_ and *x*_3_ (the values at *t* = 30 min). Those for the oscillatory regime are the limit cycle amplitude and limit cycle period (computed following section 2.13). Additionally, we allowed the biological model parameters to vary uniformly over the ranges listed in Supplemental Table A.2 with 5, 000 samples in each parameter direction. Figures 4.C and 4.D show the computed Sobol sensitivity indices for the parameters ranked by decreasing sensitivity index. Parameters that have *high* sensitivity with respect to the output quantities of interest were selected. For the bistable case, we selected the four most sensitive parameters, which were [*k*_2_, *k*_4_, *k*_5_, *k*_6_]. For the oscillatory case, we selected the parameters with a first-order sensitivity index *S*_*i*_ greater than 0.1, which were [*k*_2_, *k*_3_, *k*_4_ *k*_5_]. All remaining biological model parameters were fixed to the nominal values listed in Supplemental Table A.2. Sensitivity analysis highlighted that certain model parameters, namely *k*_2_, *k*_4_, and *k*_5_, are important in predicting both dynamical regimes. Meanwhile, parameters such as *k*_6_ and *k*_3_ are specifically important for the bistable and oscillatory regimes, respectively.

After identifiability and sensitivity analyses, we applied the CIUKF-MCMC method to estimate model parameters that predict the correct steady state. To simulate noisy experimental data, we generated two synthetic datasets (see section 2.12), sampled from the high steady state and the low steady state (circle and square markers in Fig 5.B). Each dataset had 30 full-state measurements evenly spaced over 0 *≤ t ≤* 30 (min) with measurement noise covariances set to 2.5% of the variances of the true trajectories. The prior assumptions were specified according to section 2.8 for all model parameters with the bounds listed in Supplemental Table A.2. Using the CIUKF-MCMC algorithm with AIES (section 2.9.3), we ran an ensemble of 150 Markov chains with 3, 500 samples per chain (525, 000 total samples) for each of the datasets (as shown in Supplemental Fig B.13.A-B). The maximum integrated autocorrelation times (section 2.9.5) were 190.46 for the low steady state and 169.03 for the high steady state, leading us to discard 1, 333 samples for the low steady state and 1, 183 samples for the high steady state as burn-in.

**Figure 5:**
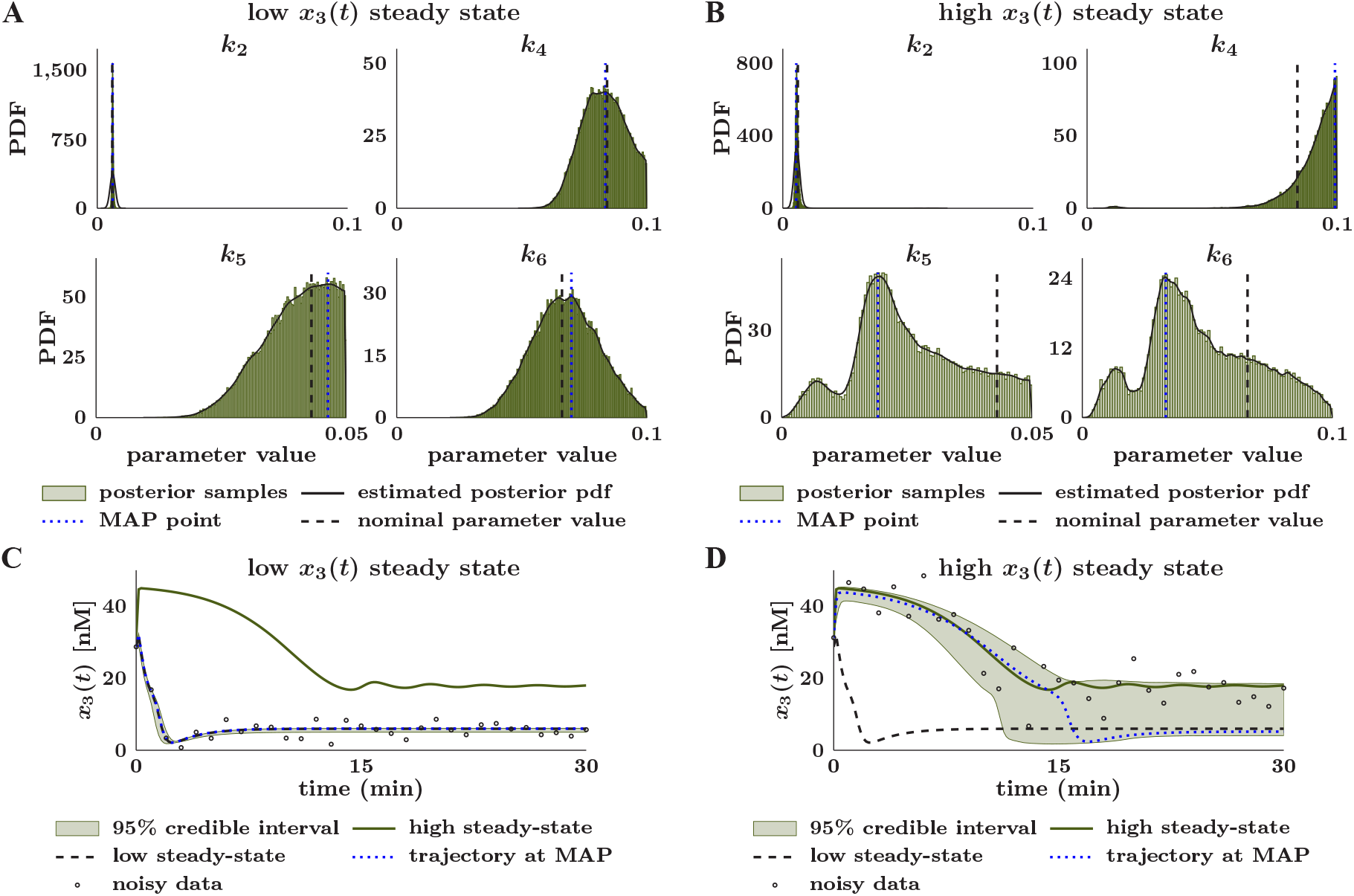
Varying levels of uncertainty in the parameters associated with the MAPK model impact steady state prediction. (**A**) Marginal posterior distributions of the model parameters for parameter estimation from noisy data of the low steady state. Posterior distributions are visualized by fitting a kernel density estimator to 325, 200 (150 walkers with 2, 167 steps each) MCMC samples obtained using CIUKF-MCMC with the affine invariant ensemble sampler (AIES) for MCMC after discarding the first 1, 333 samples per walker as burn-in. (**B**) Marginal posterior distributions of the model parameters for parameter estimation from noisy data of the high steady state reveal larger uncertainty in the model parameters when compared to the low steady state. We visualize distributions by fitting a kernel density estimator to 347, 700 (150 walkers with 2, 317 steps each) MCMC samples obtained using CIUKF-MCMC with the affine invariant ensemble sampler (AIES) for MCMC after discarding the first 1, 183 samples per walker as burn-in. (**C**) Posterior distribution of the trajectory of *x*_3_(*t*) with initial conditions that yield the low steady state highlights low uncertainty in the predicted dynamics. The true trajectory (dashed black line) shows the dynamics with the nominal parameters, the dotted blue line shows the trajectory evaluated at the MAP point, and the empty circles show the noisy data (covariance is 50% of the standard deviation of the true trajectory). The 95% credible interval shows the region between the 2.5th and 97.5th percentiles that contains 95% of 30, 000 posterior trajectories. (**D**) Posterior distribution of the trajectory of *x*_3_(*t*) with initial conditions that yield the high steady state highlights the ambiguity between which steady state is reached. All lines and computations are the same as in panel (**A**), except simulations were run using an initial condition that results in the high steady state.

The estimated marginal posterior distributions (solid black line; Fig 5.A and Fig 5.B for the low and high steady states, respectively) indicated varying levels of uncertainty between the model parameters and across the two steady states. For example, in both the low and high steady states, the marginal posterior for *k*_2_ has most of its probability mass centered around the nominal value (dashed black lines), while that for *k*_6_ has probability mass spread over a broader range of the prior support (range of the prior bounds). Additionally, the MAP points (dotted blue line) for the low steady state closely correspond to the nominal values for all model parameters, whereas there is a significant discrepancy between the MAP and the nominal values for *k*_4_, *k*_5_ and *k*_6_ in the high steady state case.

An ensemble of 30, 000 simulations with randomly selected posterior samples (see section 2.10) represented the posterior distributions of the dynamics for both steady states. For the low steady state, the trajectory evaluated at the MAP point (dotted blue line in Fig 5.C) closely matches the true trajectory (dashed black line) and the 95% credible interval (green shaded region) tightly constrains these trajectories. However, the trajectory at the MAP point for the high steady state (Fig 5.D) reaches the low steady state rather than the high steady state (solid green line). Furthermore, the 95% credible interval for the high steady state closely follows the initial transient (0 ≤ *t* ≤ 10 (min)), but it covers both steady states by the end of the simulation, e.g., for 10 ≤ *t* (min). The considerable uncertainty and bias in the estimated dynamics of the high steady state are unsurprising, given the uncertainty observed in the marginal posterior distributions. This comprehensive uncertainty analysis of the bistable MAPK dynamics showed that the presence of multiple steady states makes parameter estimation harder for the same set of parameters and governing equations. In particular the estimation uncertainty is much lower when data from the low steady state is supplied for estimation than when data from the high steady state is used.

Next, we used CIUKF-MCMC to estimate posterior distributions for the reduced set of model parameters that predict limit cycle oscillations in *x*_3_(*t*). Synthetic data with 30 samples evenly spaced over 0 ≤ *t* ≤ 90 (min) simulated noisy measurements from the oscillating trajectory at a noise level of 1% of the variance of the true trajectory (circle marks in Fig 5.B). Using CIUKF-MCMC with AIES, we ran 150 Markov chains (shown in Supplemental Fig B.14) with 6, 000 steps per chain to sample the posterior distributions. We discarded 3, 148 samples per chain, seven times the integrated autocorrelation time of 204.58, to account for burn-in. The marginal posterior distributions for [*k*_2_, *k*_3_, *k*_5_], estimated from the remaining posterior samples (Fig 6.A) show a wide range of uncertainty in the parameter estimates. For instance, the marginal posterior distributions for *k*_2_ and *k*_5_ indicate very low uncertainty, while that for *k*_4_ indicates a greater degree of uncertainty with three substantial modes in the distribution. Additionally, the distribution for *k*_3_ indicates a greater degree of uncertainty where the estimates are not constrained to one region of the domain.

**Figure 6:**
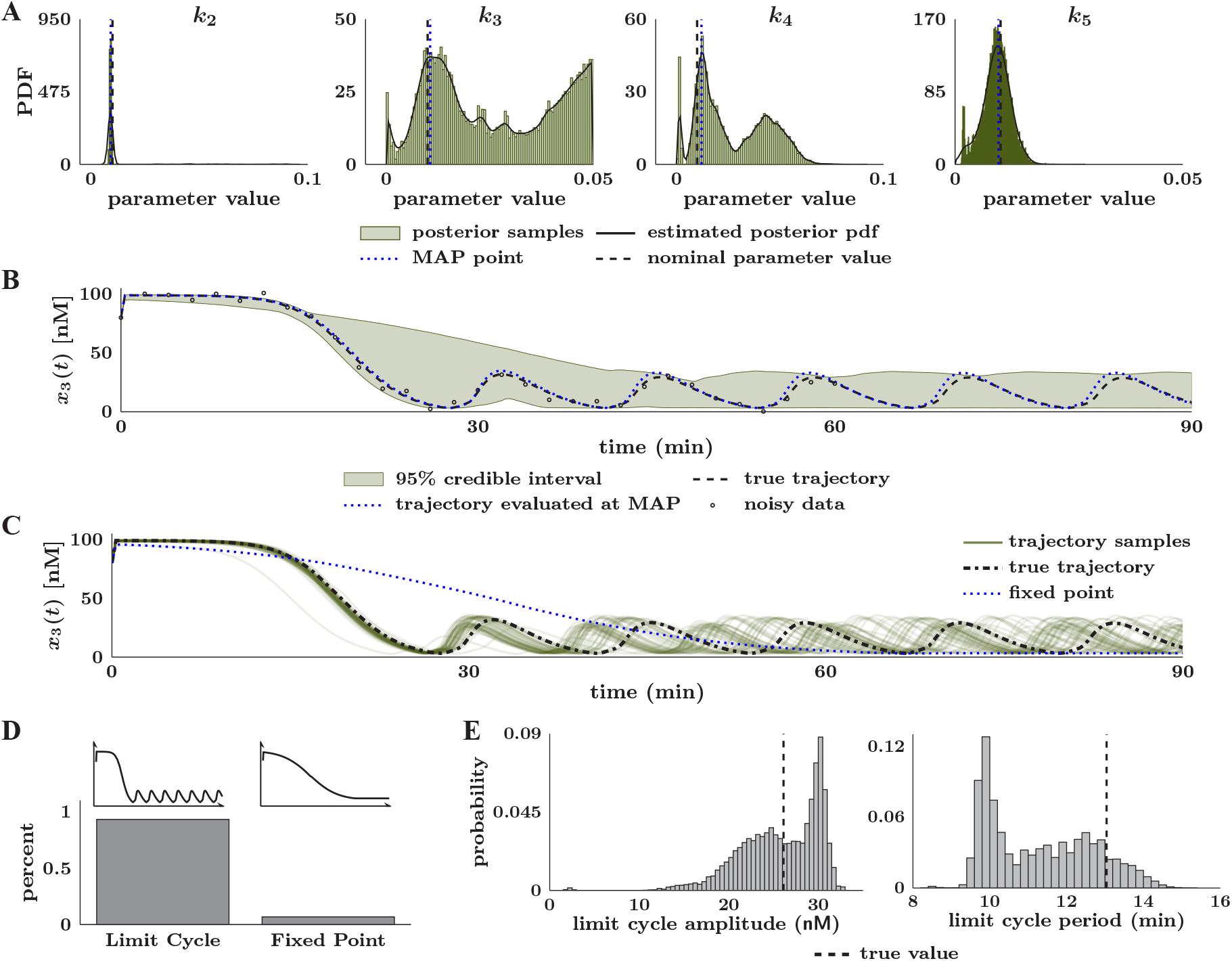
Parameter estimation results for the MAPK model in the limit cycle regime. (**A**) Marginal posterior distributions of the model parameters. Distributions are visualized by fitting a kernel density estimator to 427, 950 (150 walkers with 2, 859 steps each) MCMC samples obtained using CIUKF-MCMC with the affine invariant ensemble sampler (AIES) for MCMC after discarding the first 3, 148 samples per walker as burn-in. (**B**) Posterior distribution of the trajectory of *x*_3_(*t*) in the limit cycle regime. The true trajectory (dashed black line) shows the dynamics with the nominal parameters, the dotted blue line shows the trajectory evaluated at the MAP point, and the empty circles show the noisy data (covariance is to 1% of the variance of the true trajectory). The 95% credible interval shows the region between the 2.5th and 97.5th percentiles that contains 95% of 30, 000 posterior trajectories. (**C**) Sample posterior trajectories (50 out of 30, 000 total) reveal the variability in the limit cycle amplitude and period. Additionally, several trajectories that reach a fixed point are shown. (**D**) Quantification of the percentage of the 30, 000 sample trajectories that produce limit cycles oscillations, 93.22% (27, 966 samples), or reach a fixed point, 6.78% (2, 034 samples). (**E**) Histograms quantify the variability in limit cycle amplitude and period for the 27, 966 trajectories that show limit cycle oscillations. We define the limit cycle amplitude as the peak-to-peak difference for one oscillation, and the period is the time to complete an oscillation. The vertical blue line shows these quantities for the true trajectory.

Despite the substantial uncertainty in *k*_3_ and *k*_4_, the posterior distribution of *x*_3_(*t*) (Figure 6.B,) represented with an ensemble of 30, 000 simulations, appeared to bound the true limit cycle oscillations (see Supplemental Fig B.12.C for those of *x*_1_(*t*) and *x*_2_(*t*)). Additionally, the trajectory evaluated at the MAP point (dotted blue line) closely matches the true trajectory (dashed black line). However, the 95% credible interval (green shaded region) does not tightly constrain the dynamics as seen for the low steady state in Fig 5.C, indicating that uncertainty in *k*_3_ and *k*_4_ effects the predicted dynamics. A closer examination of a subset of 50 out of the 30, 000 posterior trajectories (green lines in Fig 6.C) revealed that the posterior trajectories included limit cycles of different amplitudes and periods along with trajectories that reach fixed points (for example, the dotted blue line in Fig 6.C). We further leveraged the posterior samples to quantify this variability. First, we separated the ensemble by the predicted dynamics, (Fig 6.D) and found that 93.22% (27, 966 samples) correctly produce limit cycle oscillations while 6.78% (2, 034 samples) reach a fixed point. Quantification of the limit cycle amplitude and period for each of the 27, 966 oscillating trajectories showed that uncertainties in the dynamics of *x*_3_(*t*) manifested in characteristics of the predicted limit cycles. Histograms of the limit cycle amplitudes and periods in Fig 6.E depict a prominent mode of amplitudes that are larger than the true amplitude and a long tail of values smaller than the true amplitude (vertical dashed black line). Additionally, there is a large mode of periods that are smaller than the period of the true trajectory. This analysis shows how uncertainty in *k*_3_ and *k*_4_ results in a range of limit cycles with different amplitudes and periods, but does not effect the ability to predict oscillatory dynamics.

Comprehensive UQ for the MAPK model highlighted how the existence of multistability introduces additional uncertainties into parameter estimation. Specifically, sensitivity analysis identified two parameters of the MAPK model, *k*_6_ and *k*_3_, that were specifically important for the bistable dynamics and the oscillatory dynamics, respectively. In both cases, Bayesian parameter estimation was able to predict the correct type of dynamics but showed remaining uncertainty in the specific characteristics of the dynamics. Overall, the proposed framework for comprehensive UQ found parameters that were important to each dynamical regime and directly quantified how uncertainties in these parameters contributes to uncertainty in the dynamics.

### 3.3. Parameter estimation in a phenomenological model of long-term potentiation/depression

A phenomenological model of coupled kinase and phosphatase switches whose activities affect the level of membrane-bound AMPAR (alpha-amino-3-hydroxy-5-methyl-4-isoxazolepropionic acid receptor) as a reporter of synaptic plasticity was proposed in [57] to capture the key events in synaptic plasticity. This kinase-phosphatase model has three states *pK*(*t*), *P* (*t*), and *A*(*t*) that correspond to active forms of kinase (CaMKII in [57]), phosphatase (PP2A in [57]), and membrane-bound AMPAR, respectively. The differential equations are

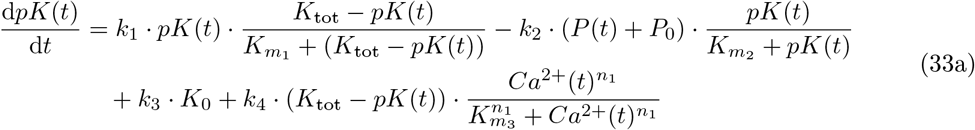

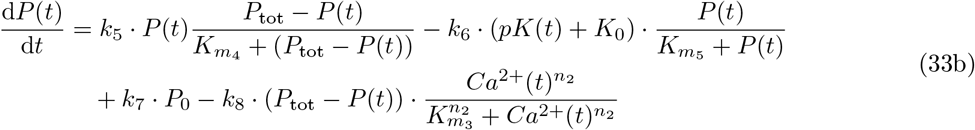

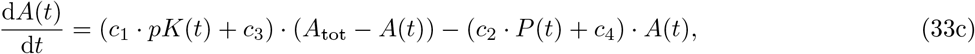

where the 24 biological model parameters are 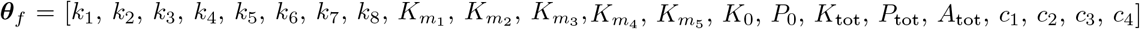. The nominal values and physiological ranges for these parameters are listed in Supplemental Table A.4.

This model predicts tristability (three steady states) in the level of excitatory postsynaptic potential (EPSP) as a function of the calcium *Ca*^2+^(*t*) input. The normalized EPSP is the membrane-bound AMPAR *A*(*t*) level, normalized to the initial condition, e.g., normalized EPSP = *A*(*t*)*/A*(*t* = 0), as defined in [57]. Fig 7.B shows simulations of the three expected responses with the nominal parameter values from [57]. The initial condition **x**_0_ = [0.0228, 0.0017, 0.4294] used for all simulations was determined by allowing the system to reach steady state with the baseline *Ca*^2+^(*t*) level of *Ca*^2+^(*t*) ≡0.1 [*µ*M]. The three steady states are the initial baseline, the higher long-term potentiation state (LTP; trajectory depicted in dashed black Fig 7.B), and the lower long-term depression state (LTD; trajectory depicted in solid green Fig 7.B). The LTP state is obtained by applying a constant stimulus of *Ca*^2+^(*t*) ≡4.0 [*µ*M] from 1 ≤ *t* ≤ 3 (sec), while the LTD state is reached by applying a constant stimulus of *Ca*^2+^(*t*) ≡ 2.2 [*µ*M] in the same time interval; *Ca*^2+^(*t*) is set to the baseline level before (*t <* 1 (sec)) and after (*t >* 3 (sec)) the stimulus is applied. We investigated how well the proposed uncertainty quantification framework could estimate the model parameters for LTP and LTD from synthetic data of an LTP-inducing calcium input.

**Figure 7:**
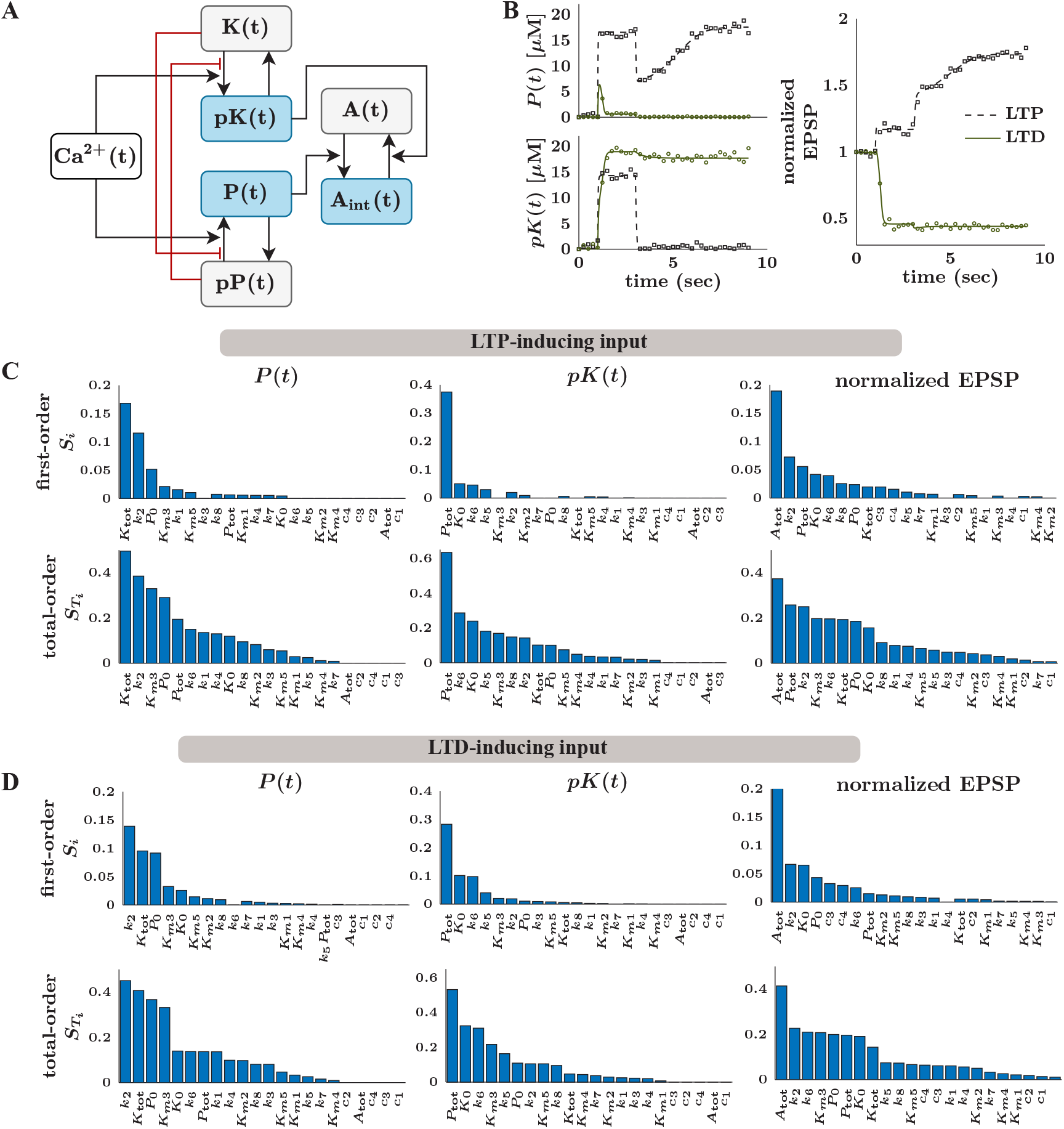
Parameter estimation for a coupled kinase-phosphatase switch for long-term potentiation and long-term depression in neurons as a function of calcium input. (**A**) Network diagram of the simplified coupled kinase-phosphatase signaling model where calcium *Ca*^2+^(*t*) acts as the input. (**B**) Trajectories of the three state variables in response to long-term potentiation (LTP; pulse of *Ca*^2+^(*t*) ≡ 4.0 [*µ*M] from 2 ≤ *t* ≤ 3 (sec)) and long-term depression (LTD; pulse of *Ca*^2+^(*t*) ≡ 2.2 [*µ*M] from 2 ≤ *t* ≤ 3(sec)) inducing calcium inputs. The calcium level is set to a baseline of *Ca*^2+^(*t*) ≡ 0.1 [*µ*M] before and after stimulus. We compute normalized EPSP by normalizing *A*(*t*) to its initial condition as described in [57]. The synthetic noisy data for the LTP and LTD cases are indicated by the black square and green circle marks, respectively, with the noise covariance equal to 1% of the variance of the data. (**C** and **D**) Sobol sensitivity indices for all free model parameters in response to LTP-inducing and LTD-inducing inputs, respectively. The quantities of interest are the steady state values of each state variable. We show both the first-order sensitivity indices *S*_*i*_ and the total-order indices 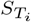. We select a reduced set of free parameters by choosing the parameters whose first-order sensitivity index is greater than 0.05, e.g., *S*_*i*_ *>* 0.05. This gives us the same set of free parameters, ***θ***_*f*_ = [*k*_2_, *k*_6_, *K*_0_, *P*_0_, *K*_tot_, *P*_tot_, *A*_tot_], for both the LTP and LTD cases. Remaining model parameters are fixed to the nominal values in Supplemental Table A.4.

Following the proposed framework, the parameter space is reduced by performing identifiability and sensitivity analysis. First, identifiability analysis showed that all model parameters, except *n*_1_ and *n*_2_, are globally structurally identifiable from full-state measurements. Next, global sensitivity analysis of the steady states in response to an LTP-inducing and an LTD-inducing input ranked the 22 globally identifiable model parameters. All free parameters were uniformly varied over two orders of magnitude centered around the nominal values, 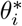, for each parameter, e.g. 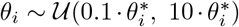. In order to maintain conservation of mass, the lower bounds of the total concentration parameters, *K*_tot_, *P*_tot_, *A*_tot_, were chosen to be greater than or equal to the initial condition. Figure 7.C shows the computed Sobol sensitivity indices for the LTP-inducing input and Fig 7.D shows those for the LTD-inducing input. The sensitivity analyses point to the same group of seven model parameters for both the LTP-inducing and LTD-inducing inputs. These are ***θ***_*f*_ = [*k*_2_, *k*_6_, *K*_0_, *P*_0_, *K*_tot_, *P*_tot_, *A*_tot_], whose first-order indices were greater than 0.05, e.g., *S*_*i*_ *>* 0.05. We chose to estimate these seven parameters and fix the remaining model parameters to the nominal values listed in Supplemental Table A.4.

Next, we used the CIUKF-MCMC algorithm with AIES to estimate the posterior distribution for the reduced set of parameters. The parameters were estimated from noisy synthetic data with 36 full-state measurements of the LTP response. The data are spread uniformly over the domain 0 ≤ *t* ≤ 9 (sec) at a noise level of 1% of the variance of the true trajectory for the respective states. The maximum integrated autocorrelation time of the ensemble 150 Markov chains with 8, 000 steps per chain was 621.58, leading us to discard 4, 351 samples as burn-in. Traces of the ensemble of Markov chains for all parameters are shown in Supplemental Fig B.15. Figure 8.A shows the estimated marginal posterior distributions for the free model parameters, *k*_2_, *k*_6_, *P*_0_, *K*_0_, *P*_tot_, *K*_tot_, *A*_tot_. We observed different levels of uncertainty across the estimated parameters. For instance, the marginal posterior for *A*_tot_ indicated a very high level of certainty with almost all of the probability mass, and thus the MAP point, aligned to the nominal value. However, the marginal posterior distributions for *P*_0_ and *K*_0_ show much more significant uncertainty because we observe posterior probability spread over the entire support of the prior. Additionally, the marginal posterior distributions for *k*_2_, *k*_6_, *P*_tot_, and *K*_tot_ show a large mode around the MAP point that is shifted from the nominal value and a smaller mode at the nominal value.

**Figure 8:**
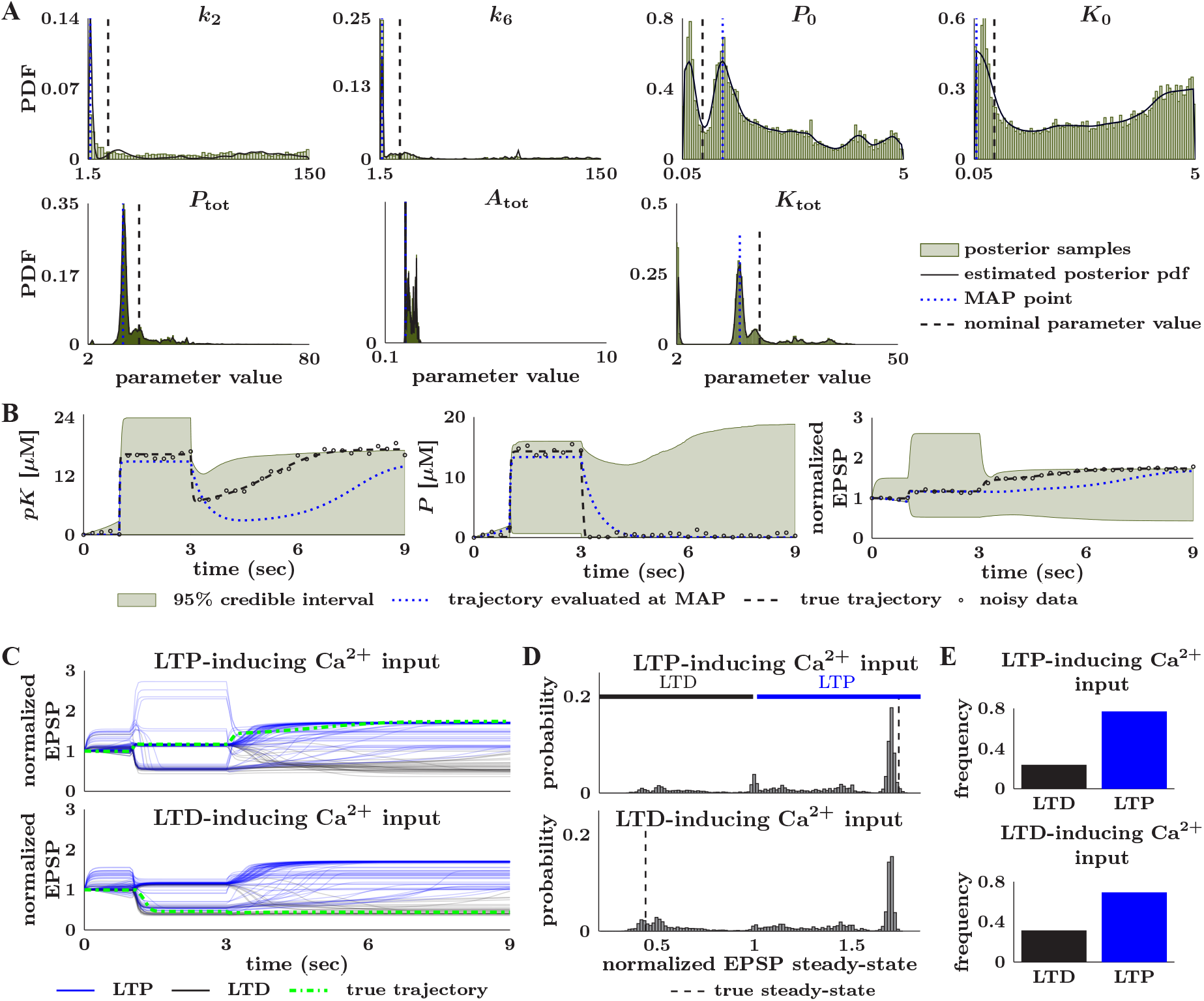
Comprehensive parameter estimation and uncertainty quantification reveal failures to predict the correct long-term model behavior. (**A**) We estimated marginal posterior distributions of the model parameters from noisy data with an LTP-inducing calcium input. Distributions are visualized by fitting a kernel density estimator to 547, 500 (150 walkers with 3, 649 steps each) MCMC samples obtained using CIUKF-MCMC with AIES for MCMC after discarding the first 4, 351 samples per walker as burn-in. (**B**) Posterior distribution of the trajectories of the state variables show LTP (normalized EPSP *>* 1) and LTD (normalized EPSP *<* 1) responses for an LTP inducing input. The true trajectory (dashed black line) shows the dynamics with the nominal parameters, the dotted blue line shows the trajectory evaluated at the MAP point, and the points show the noisy data (covariance is 1% of the variance of the true trajectory). The 95% credible interval shows the region between the 2.5th and 97.5th percentiles that contains 95% of 30, 000 posterior trajectories. (**C**) Sample posterior trajectories (100 out of 30, 000 total) highlight the LTP (blue lines) and LTD (black lines) in response to the LTP inducing input (top) and the LTD inducing input (bottom). The dashed green lines show the true trajectories for the respective calcium inputs. (**D**) Histograms reveal the distribution of the long-term responses to the LTP-inducing input (top) and the LTD-inducing input (bottom). The dashed black lines show the true response for the respective calcium inputs. (**E**) Quantifying the percentage of the 30, 000 sample trajectories that produce LTD and LTP responses for each calcium input. The LTP-inducing input yields 76.59% (22, 972 samples) of the responses in the LTP state and 23.41% (7, 024 samples) in the LTD state. The LTD inducing input yields 68.93% (20, 680 samples) of the responses in the LTP state and 31.07% (9, 320 samples) in the LTD state.

Using the posterior samples, an ensemble of 30, 000 simulations (see section 2.10) represented the posterior distribution of the predicted dynamics in response to an LTP-inducing input, as shown in Fig 8.B. We observed that the 95% credible interval for the normalized EPSP covered both LTP (normalized EPSP *>* 1) and LTD (normalized EPSP *<* 1) responses even though the input was LTP-inducing. However, the trajectory evaluated at the MAP point (dotted blue line) matched the true trajectory (dashed black line), indicating that most trajectories align with the expected LTP response. Examination of the individual trajectories within the ensemble simulation (Fig 8.C, top row) and the normalized EPSP steady state values (Fig 8.D, top row) confirmed that there are both LTP (blue traces) and LTD (black traces) responses to the LTP-inducing input. Specifically, we found that 76.59% (22, 972 samples) of the responses correctly predict LTP, and 23.41% (7, 024 samples) of response incorrectly predict LTD (Fig 8.E). Therefore, despite the high-quality data (many measurements and low noise), we still observed substantial uncertainty in the predicted normalized EPSP.

Lastly, we investigated if the posterior distribution estimated in the LTP regime can predict the response to an LTD-inducing input. An initial hypothesis was that in response to an LTD-inducing input, most trajectories would predict LTD, but a significant subset would predict the incorrect response, LTP. An additional ensemble of 30, 000 simulations, with an LTD-inducing input of *Ca*^2+^(*t*) ≡ 2.2 [*µ*M] (pulse from 2 ≤ *t* ≤ 3 (sec)) was used to determine the posterior distribution of the dynamics. Visualization of 100 of the 30, 000 trajectories in Fig 8.C (bottom row) again showed both LTP (blue) and LTD (black) responses; however, we unexpectedly observed more LTP than LTD in response to the LTD-inducing input. The distribution of normalized EPSP responses (bottom row of Fig 8.D) confirms that there are a large number of responses in the LTP regime with a minor mode around the correct response. Quantification of these results highlights that only 31.07% (9, 320 samples) of responses are in the LTD state (expected response) while 68.93% (20, 680 samples) of the responses are in the LTP state (unexpected response). In summary, this model formulation can correctly capture the same LTP behavior over a range of model parameters while losing the ability to predict LTD behavior.

Despite the LTP and LTD responses being sensitive to the same set of parameters (see Fig 7.C-D), the posterior distribution estimated from measurements of LTP places more probability on parameter sets that robustly predict LTP over those that correctly predict both responses. From these results, we conclude that for this model, we need to learn the model parameters with a high degree of certainty in order to disambiguate the LTP versus the LTD response because sensitivity analyses revealed that the same set of parameters governs these two different outputs. This finding highlights that sensitivity analyses are not sufficient to distinguish parameter uncertainty for systems with multistability and a comprehensive framework as outlined here is necessary to shed light on such model complexities.

## 4. Discussion

In this work, we developed a framework (see section 2.1) for comprehensive uncertainty quantification of dynamical models in systems biology. The proposed framework leverages identifiability and sensitivity analysis to reduce the parameter space (sections 2.3 and 2.4) followed by Bayesian parameter estimation with CIUKF-MCMC (see section 2.6). We applied this framework to three systems biology models to demonstrate its applicability and highlight how a focus on uncertainty can transform modeling-based studies. First, we performed two computational experiments on a simple two-state model that showed how noise and data sparsity contribute to estimation uncertainty. Next, we applied our framework to two models, the MAPK and the synaptic plasticity models, which better resemble the models used to capture biological readouts. Using these models, we highlight how comprehensive uncertainty quantification enables quantitative analysis of two biologically relevant dynamical behaviors, limit cycles and steady state responses. We also found that good quality data cannot always overcome uncertainty due to the model structure. These examples provide an essential starting point for applying our framework in practice and interpreting systems biology studies under uncertainty.

Our results highlight how a focus on uncertainty quantification can give new insights in modeling-based studies. For example, in section 3.2 we were able to learn a posterior distribution for the parameters that predict limit cycles over a range of amplitudes and periods. The posterior distribution is our *best guess* for the distribution of the model parameters after incorporating the available data into the statistical model. Therefore, the posterior distribution for the dynamics (approximated via an ensemble simulation as in section 2.10) provides our *best guess* for the dynamics, given everything we know about the model. For the MAPK limit cycle oscillations, this *best guess* includes dynamics with a range of limit cycle properties. As highlighted in section 3.1, we can expect the quality of this guess to reflect the quality and quantity of the available experimental data. Therefore, incorporating uncertainty quantification into modeling provides additional context for interpreting the predictions. We also observe that predictions do not always capture the correct bistability in the example of high MAPK steady state (section 3.2) and the LTP/LTD response (section 3.3). In both cases, the 95% credible intervals of the ensemble of predicted dynamics cover both the higher and lower steady states (LTP and LTD in the synaptic plasticity example). These results imply that the estimated posterior distributions for these models include parameter sets that no longer show bistability at the specified input (the initial condition for the MAPK model or the calcium level for the synaptic plasticity model). Further analysis of these systems could test if these parameter sets lead to a complete loss of bistability or merely shift the bifurcation point, the value of the input that changes which steady state is reached. Overall our results point to a complex interplay between model parameters and inputs that potentially confounds parameter estimation of multistable systems.

Throughout this work, we assume that the model equations are known prior to parameter estimation. This assumption reflects standard modeling practices where models use biochemical theories that assume equations for the kinetics of biochemical reactions. In using the CIUKF-MCMC we somewhat weaken this assumption because it introduces process noise to account for uncertainty in the model form [26]. In reality, all models have some level of uncertainty because they rely on assumptions about the system. Therefore, accounting for model form uncertainty regularizes the dynamics to account for a mismatch between the predicted dynamics and the data [26]. However, it may be necessary to simultaneously estimate the model structure (formulation of the equations) and parameters from the data. One approach to disambiguate a model structure is to learn the biochemical reaction network [91, 92] or the mathematical model directly from data [93, 94, 95, 96, 97]. Additionally, it is possible to cast these problems in the Bayesian perspective to learn the model form and the associated uncertainties [26, 98].

While Bayesian methods are well suited for uncertainty quantification in systems biology, it is also important to understand the caveats associated with MCMC methods (see section 2.9). Markov chain Monte Carlo sampling is inherently compute-intensive [19, 27, 77] and requires careful analysis to apply correctly [77]. Most importantly, there is always a possibility that the distribution of the samples may not have completely converged to the true posterior despite indications that it has [27, 76, 77]. In this work, we take several steps, computing the integrated autocorrelation time and visualizing the Markov chains, to mitigate these problems and improve our confidence in the MCMC results. Different approaches to assessing Markov chains are presented in detail in [27, 76, 77]. Additionally, the affine invariant ensemble sampler is only well suited to sample anisotropic posterior distributions in small to moderate parameter spaces, up to about 50 parameters [59, 99]. In this work, we focus on models with at most three state variables and no greater than 22 model parameters, but many systems biology models can have 10s of states with 10s-100s of parameters [13, 28]. While we showed that identifiability and sensitivity analyses could significantly reduce the number of free parameters, how these conclusions might change for larger systems remains to be tested. Potential approaches for sampling such spaces include variational inference [100] and randomize then optimize approaches [101].

Lastly, we make several assumptions in choosing statistical models for measurement and process noise and in simulating biological measurement data. First, we assume Gaussian measurement and process noises; however, we may be better able to describe noise in biological systems and experiments with alternate probability distributions (see e.g., [46]). Assuming a non-Gaussian distribution for the measurement noise would require reformulating the likelihood function (it would alter the distribution in Eq (12)); however, incorporating alternative distributions for the process noise would require significant effort. The constrained interval unscented Kalman filter (also most other Kalman filters) revolves around the assumption of normally distributed process noise [68]; thus, alternative noise models would require approximations for the marginal likelihood that go beyond Kalman filtering. Next, this work considers linear measurement functions, see e.g., Eq (3), but CIUKF-MCMC is also well-suited to handle nonlinear measurement functions [26]. Lastly, we assume that we only have access to a single time series of measurements, e.g., one trial of an experiment; however, most experiments in biology perform several repeated trials. In these cases, one would like to incorporate all available data to inform the statistical model. A straightforward approach would use the mean of each time point and estimate parameters from the time series of means. Additionally, one could estimate parameters from each time series separately and then analyze several posterior distributions. These separate distributions can then be merged via meta-analysis or information fusion principles [102, 103, 104]. Lastly, one could construct a statistical model that accounts for the multiple time series [27] simultaneously. In summary, research at the intersection of uncertainty quantification and systems biology modeling will strengthen parameter estimation and enable models that more accurately represent experimental measurements.

## 5. Code availability statement

The code necessary to reproduce this work is available on GitHub at https://github.com/RangamaniLabUCSD/CIUKF-MCMC.

## 6. Acknowledgements

The authors would like to thank Miriam Bell, Sage Malingen, María Hernández Mesa, Lingxia Qiao, Mayte Bonilla Quintana, and other members of the Rangamani and Kramer groups for critical discussions and feedback on the manuscript. Additionaly the authors would like to thank Matthias Morzfeld, Alex Gorodetsky and Nick Galioto for their methodological insights and helpful discussions. Nathaniel Linden acknowledges support from the National Institute of Biomedical Imaging and Bioengineering (NIBIB) of the National Institutes of Health (NIH) under award number T32EB9380 and a UCSD Sloan Scholar Fellowship. Padmini Rangamani acknowledges support from Air Force Office of Scientific Research (AFOSR) Multidisciplinary University Research Initiative (MURI) grant FA9550-18-1-0051.

## Appendix A. Supplemental Tables

This section provides the tables that list initial conditions, nominal parameter values, and relevant parameter ranges for the models presented in section 3. Table A.1 corresponds to the two state model in section 3.1. Tables A.2 and A.3 correspond to the MAPK model in section 3.2. Lastly, table A.4 corresponds to the synaptic plasticity model in section 3.3.

**Table A.1:**
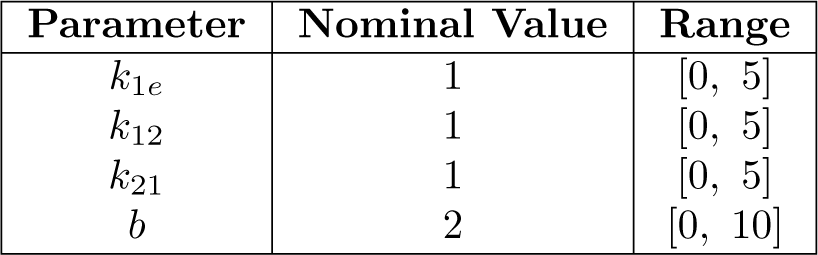
Two-compartment model [55] model parameters and relevant ranges. All listed values have units of one over time.

**Table A.2:** MAPK model parameters and relevant ranges from [56]. Note: For the oscillatory dynamics, the range for *k*_5_ is [1 *×* 10^*−* 5^ 0.05].

| Parameter | Nominal Value: Limit-cycle | Nominal Value: Bistable | Range |
| --- | --- | --- | --- |
| $S_{1t}$ | 100 nM | 0.22 nM | [0, 100] |
| $S_{2t}$ | 100 nM | 10 nM | [0, 100] |
| $S_{3t}$ | 100 nM | 53 nM | [0, 100] |
| $k_1$ | $0.1 \text{ nM} \cdot \text{s}^{-1}$ | $0.0012 \text{ nM} \cdot \text{s}^{-1}$ | [0, 0.05] |
| $k_2$ | $0.01 \text{ nM} \cdot \text{s}^{-1}$ | $0.006 \text{ nM} \cdot \text{s}^{-1}$ | [0, 0.1] |
| $k_3$ | $0.01 \text{ nM} \cdot \text{s}^{-1}$ | $0.049 \text{ nM} \cdot \text{s}^{-1}$ | [0, 0.05] |
| $k_4$ | $0.01 \text{ nM} \cdot \text{s}^{-1}$ | $0.084 \text{ nM} \cdot \text{s}^{-1}$ | [0, 0.1] |
| $k_5$ | $0.01 \text{ nM} \cdot \text{s}^{-1}$ | $0.043 \text{ nM} \cdot \text{s}^{-1}$ | [0, 0.05] |
| $k_6$ | $0.01 \text{ nM} \cdot \text{s}^{-1}$ | $0.066 \text{ nM} \cdot \text{s}^{-1}$ | [0, 0.1] |
| $n_1$ | 10 | 5 | [5, 10] |
| $K_1$ | 1 nM | 9.5 nM | [0, 10] |
| $n_2$ | 15 | 10 | [5, 10] |
| $K_2$ | 8 nM | 15 nM | [0, 20] |
| $\alpha$ | 10 | 95 | [0, 100] |

**Table A.3:** Initial conditions used for simulations of the MAPK model from [56].

| State-variable | Limit cycle | Bistable: low steady state | Bistable: high steady state |
| --- | --- | --- | --- |
| $x_1(0)$ | 10 nM | 0.0015 nM | 0.1245 nM |
| $x_2(0)$ | 80 nM | 3.6678 nM | 2.4870 nM |
| $x_3(0)$ | 80 nM | 28.7307 nM | 31.2623 nM |

**Table A.4:** Long-term potentiation/depression model parameters from [57] and ranges. The ranges are given by [0.1 · 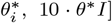, where 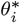 is the nominal value. Note: we do not include ranges for *n*_1_ and *n*_2_ because these parameters are always set to the nominal values.

| Parameter | Nominal Value | Range |
| --- | --- | --- |
| $k_1$ | $2 \text{ s}^{-1}$ | $[0.2, 20]$ |
| $k_2$ | $15 \text{ s}^{-1}$ | $[1.5, 150]$ |
| $k_3$ | $1 \text{ s}^{-1}$ | $[0.1, 100]$ |
| $k_4$ | $120 \text{ s}^{-1}$ | $[12, 1200]$ |
| $k_5$ | $2 \text{ s}^{-1}$ | $[0.2, 20]$ |
| $k_6$ | $15 \text{ s}^{-1}$ | $[1.5, 150]$ |
| $k_7$ | $1 \text{ s}^{-1}$ | $[0.1, 10]$ |
| $k_8$ | $80 \text{ s}^{-1}$ | $[8.0, 800]$ |
| $c_1$ | 1 | $[0.1, 10]$ |
| $c_2$ | 1 | $[0.1, 10]$ |
| $c_3$ | $6 \text{ s}^{-1}$ | $[0.6, 60]$ |
| $c_4$ | $8 \text{ s}^{-1}$ | $[0.8, 80]$ |
| $K_{m1}$ | $10 \text{ }\mu\text{M}$ | $[1.0, 100]$ |
| $K_{m2}$ | $0.3 \text{ }\mu\text{M}$ | $[0.03, 3]$ |
| $K_{m3}$ | $4 \text{ }\mu\text{M}$ | $[0.4, 40]$ |
| $K_{m4}$ | $10 \text{ }\mu\text{M}$ | $[1.0, 100]$ |
| $K_{m5}$ | $1 \text{ }\mu\text{M}$ | $[0.1, 10]$ |
| $K_0$ | $0.5 \text{ }\mu\text{M}$ | $[0.05, 5]$ |
| $P_0$ | $0.5 \text{ }\mu\text{M}$ | $[0.05, 5]$ |
| $K_{\text{tot}}$ | $20 \text{ }\mu\text{M}$ | $[2, 200]$ |
| $P_{\text{tot}}$ | $20 \text{ }\mu\text{M}$ | $[2, 200]$ |
| $A_{\text{tot}}$ | 1 | $[0.1, 10]$ |
| $n_1$ | 4 | — |
| $n_2$ | 3 | — |

## Appendix B. Supplemental Figures

This section provides supplemental figures for the results presented in section 3. Figures B.9, B.10, and B.11 correspond to section 3.1. Figures B.12, B.13, and B.14 correspond to section 3.2. Lastly, Fig B.15 corresponds to section 3.3.

**Figure B.9:**
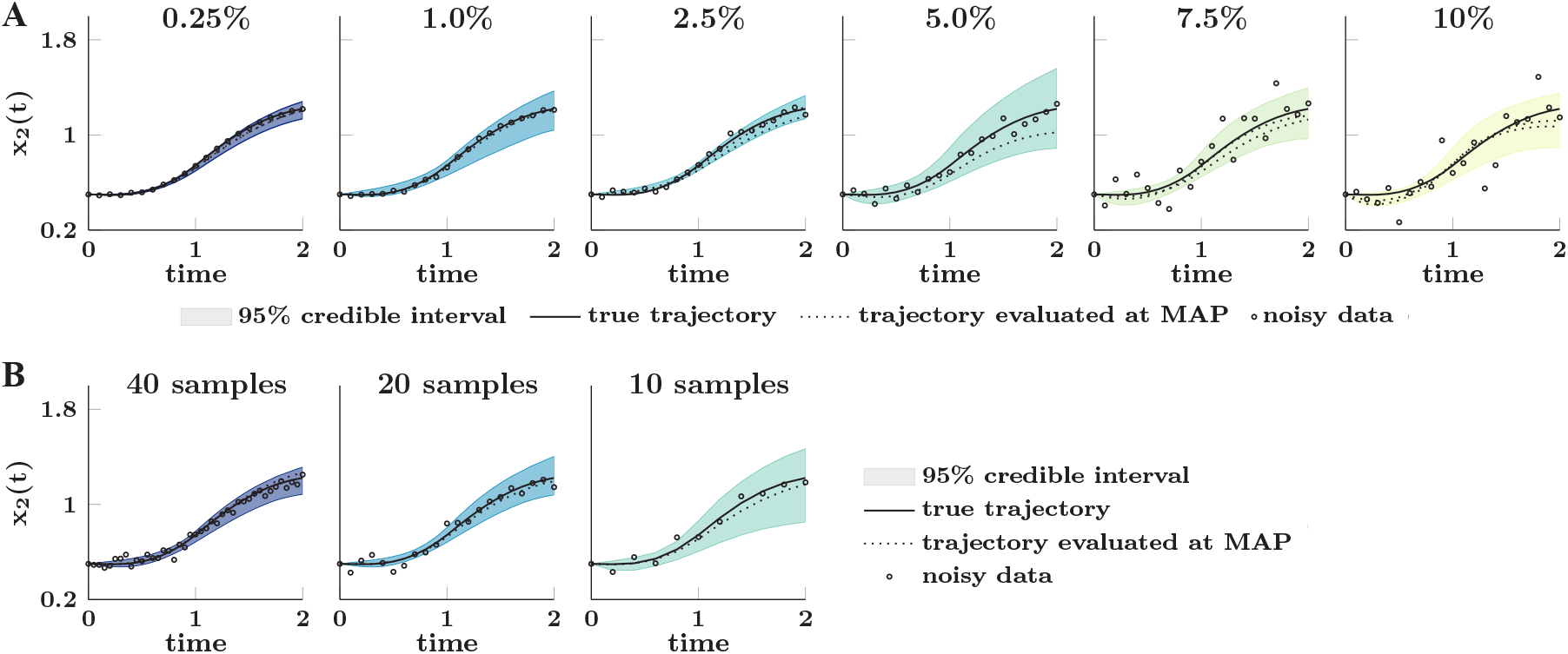
Posterior distributions of *x*_2_(*t*) in the two-state model experiments. (**A**) corresponds to the measurement noise experiment, Fig 3.C. We observe that increasing the data noise level increased the uncertainty in the predicted dynamics. We control the noise level by setting the noise covariances to the specified percentage of the standard deviation of each state variable. The dashed black vertical lines indicate each parameter’s nominal (true) value. (**B**) corresponds to the data sparsity experiment, Fig 3.E. We observe that decreasing the number of experimental samples supplied for estimation increased estimation uncertainty. The noise level in the data was fixed to the 2.5% level shown above

**Figure B.10:**
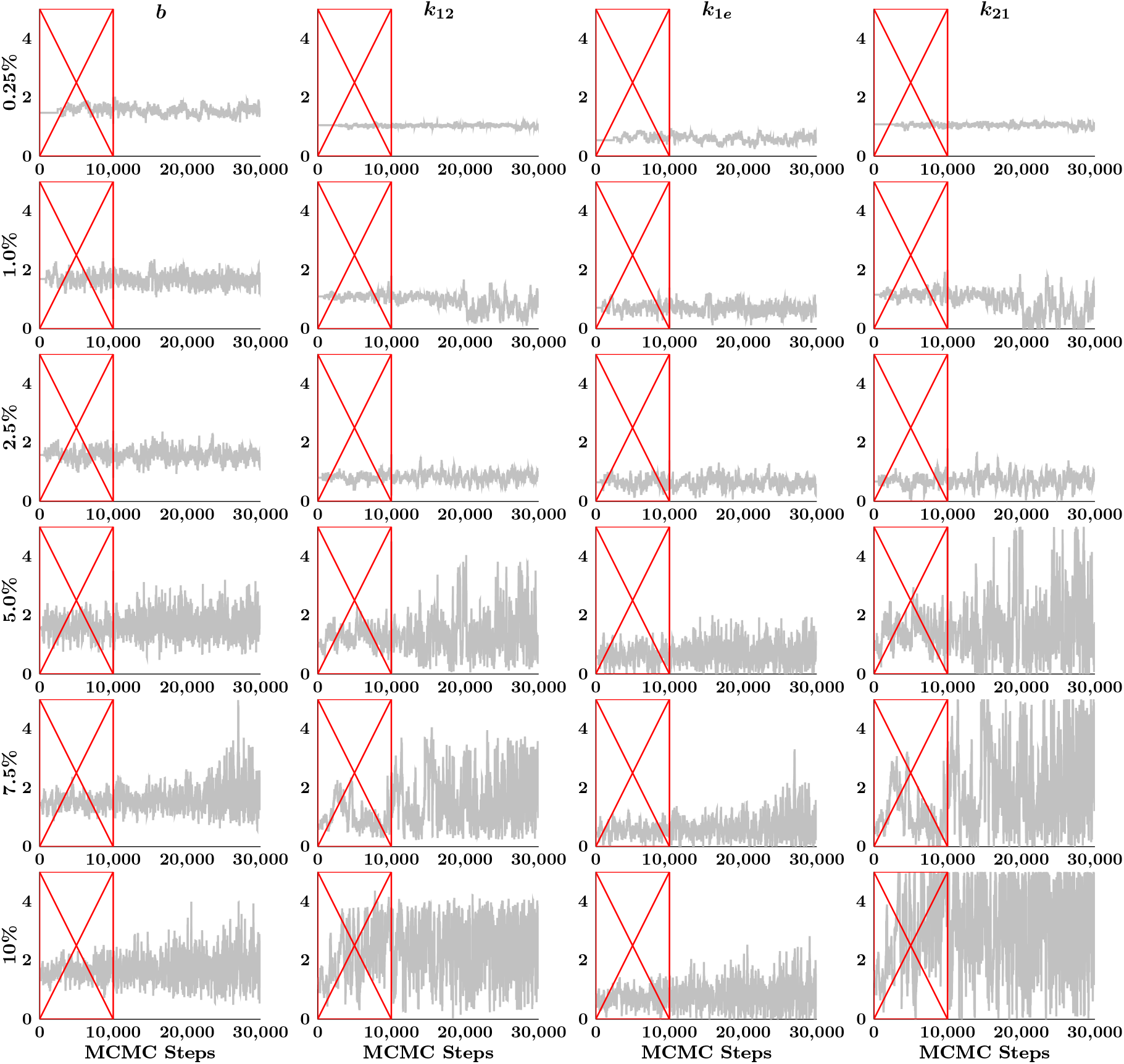
Markov chains for the two-state model parameters at increasing noise levels in the measurement noise experiment, Fig 3.B-C. Each row corresponds to a noise level; noise increases down the figure. The red boxes indicate samples discarded as burn-in.

**Figure B.11:**
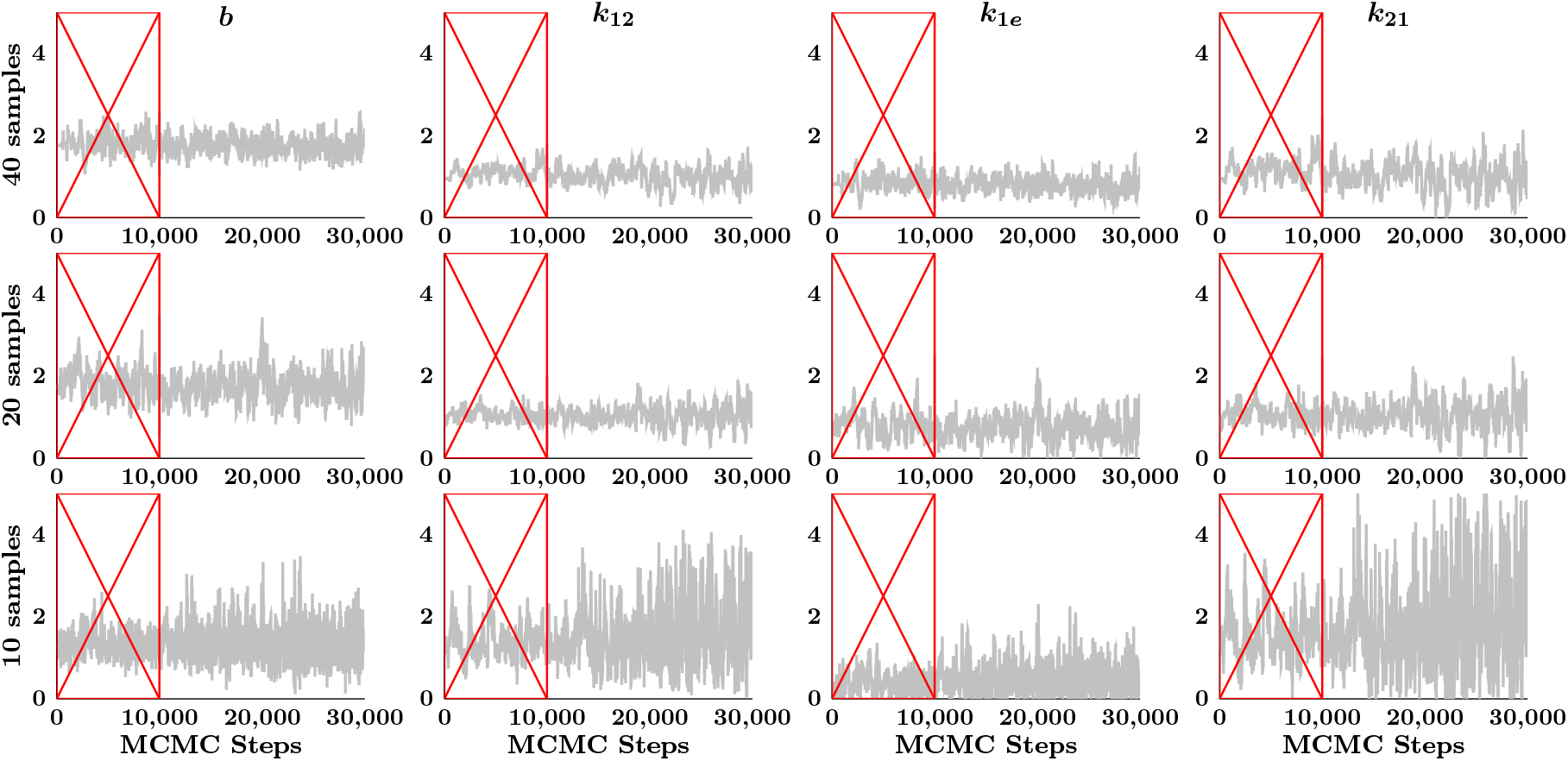
Markov chains for the two-state model parameters with decreasing samples in the measurement data sparsity experiment, Fig 3.D-E. Each row corresponds to a sparsity level. The red boxes indicate samples discarded as burn-in.

**Figure B.12:**
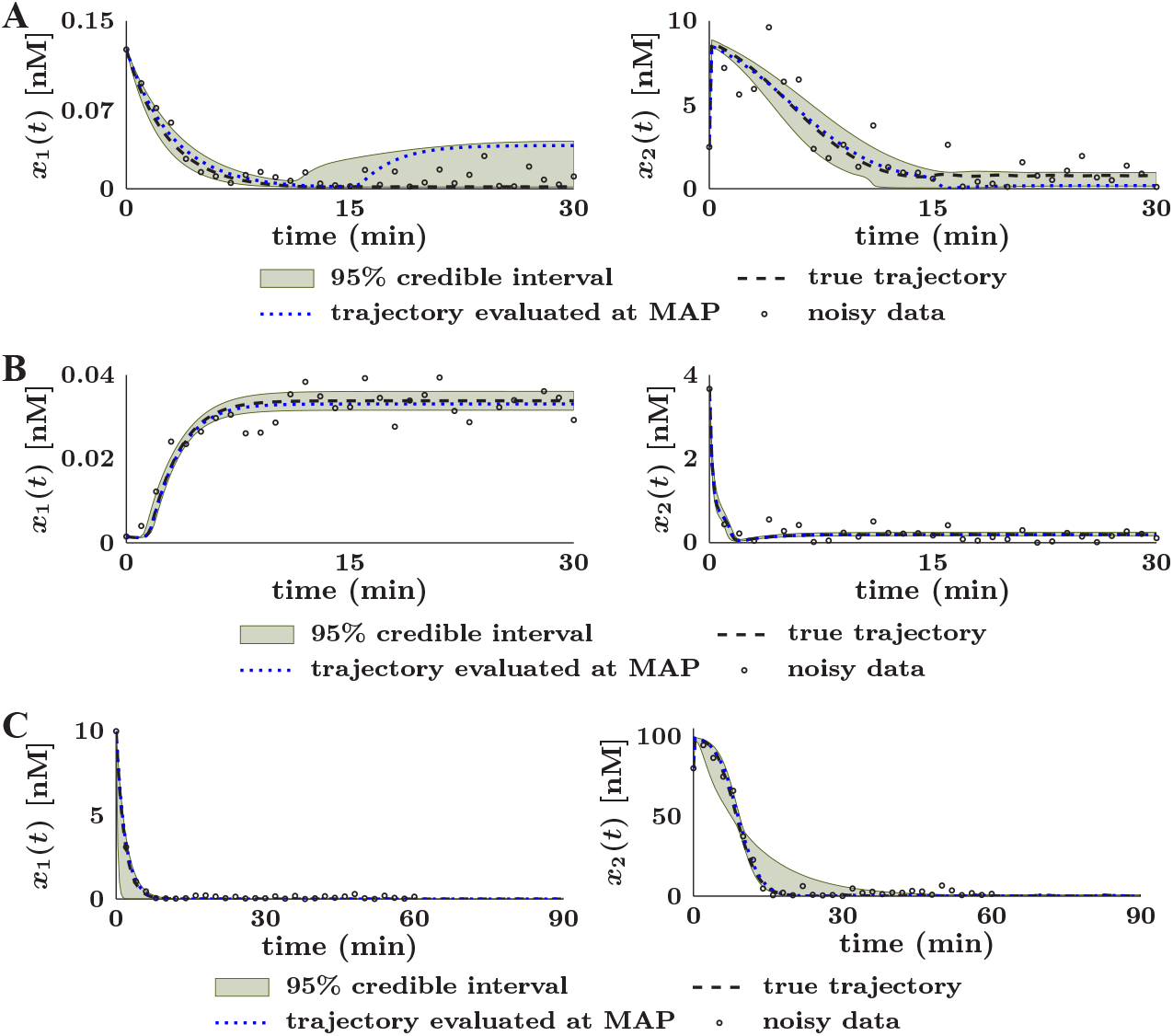
Posterior distributions of *x*_1_(*t*) and *x*_2_(*t*) for the MAPK. (**A**) Trajectories for the high steady-state; corresponds to Fig 5.D. (**A**) Trajectories for the low steady-state; corresponds to Fig 5.C. (**B**) Trajectories for limit cycle oscillations; corresponds to Fig 6.B.

**Figure B.13:**
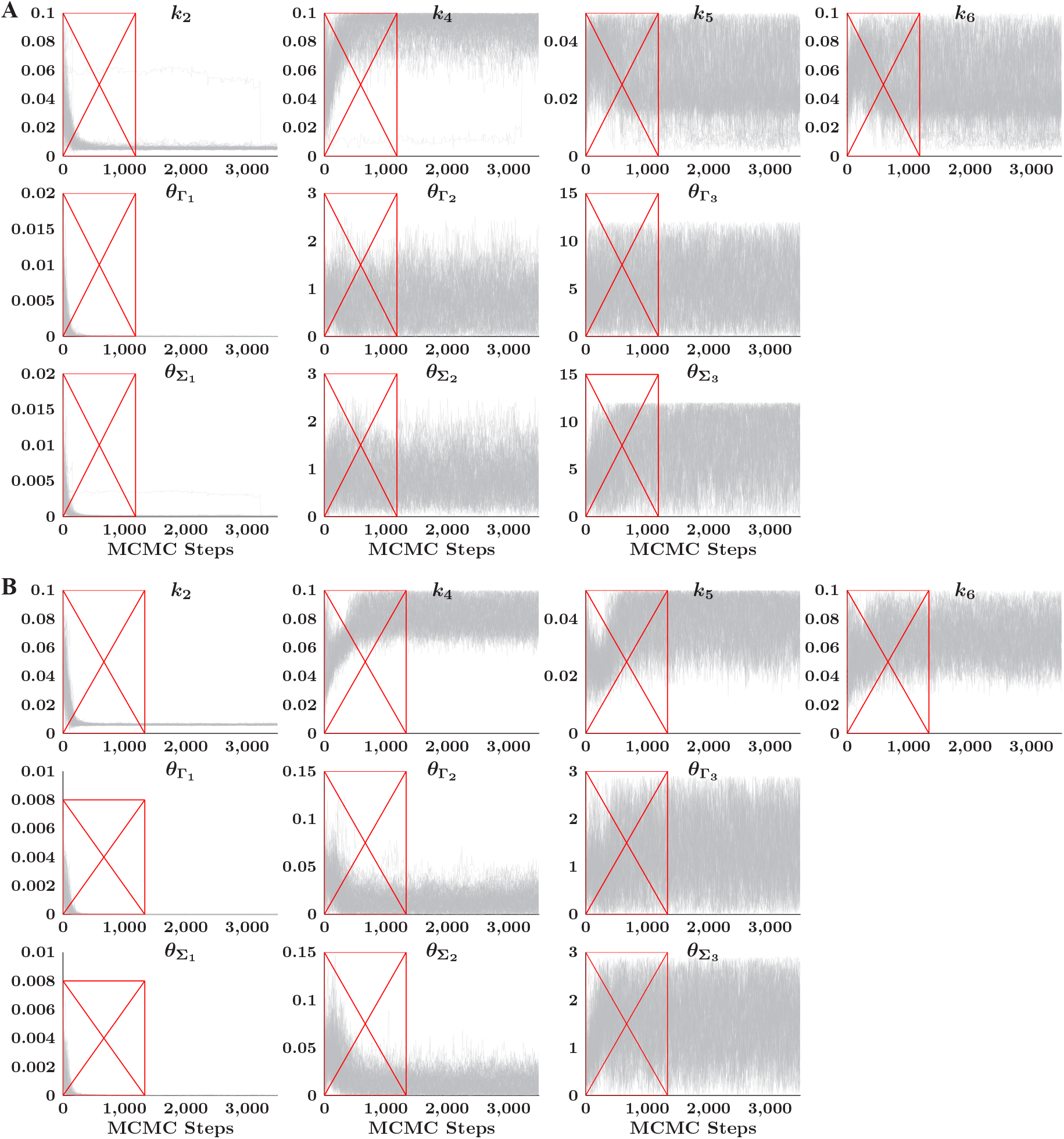
Markov chains for the MAPK model parameters and UKF-MCMC noise covariances for bistability. (**A**) Low steady-state, Fig 5.A. (**B**) High steady-state, Fig 5.B. The process noise covariances are 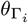 and 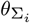 are the measurement noise covariances. The red boxes indicate samples discarded as burn-in.

**Figure B.14:**
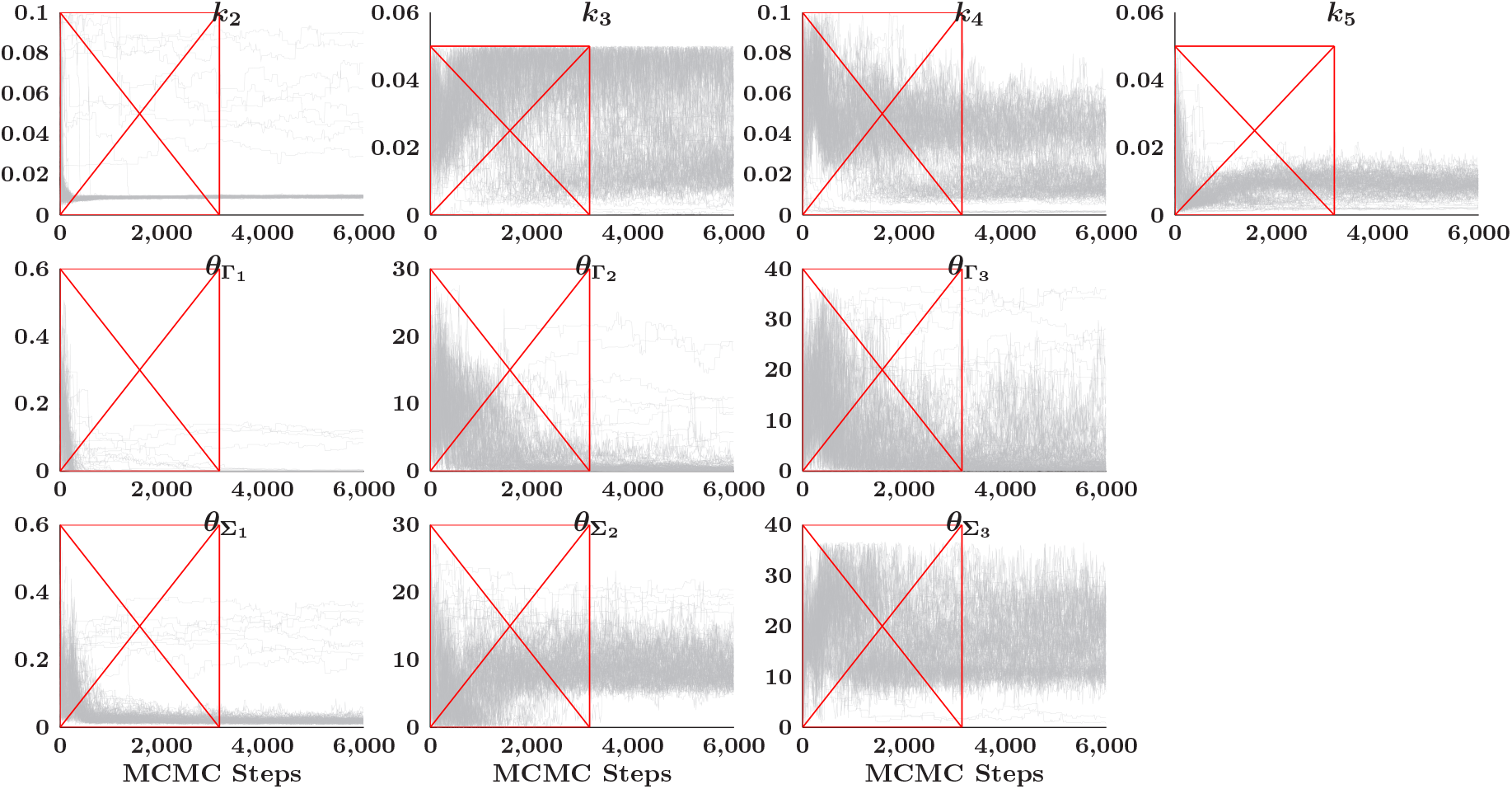
Markov chains for the MAPK model parameters and UKF-MCMC noise covariances for limit cycle oscillations; corresponds to 6.A The process noise covariances are 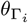 and 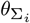 are the measurement noise covariances. The red boxes indicate samples discarded as burn-in.

**Figure B.15:**
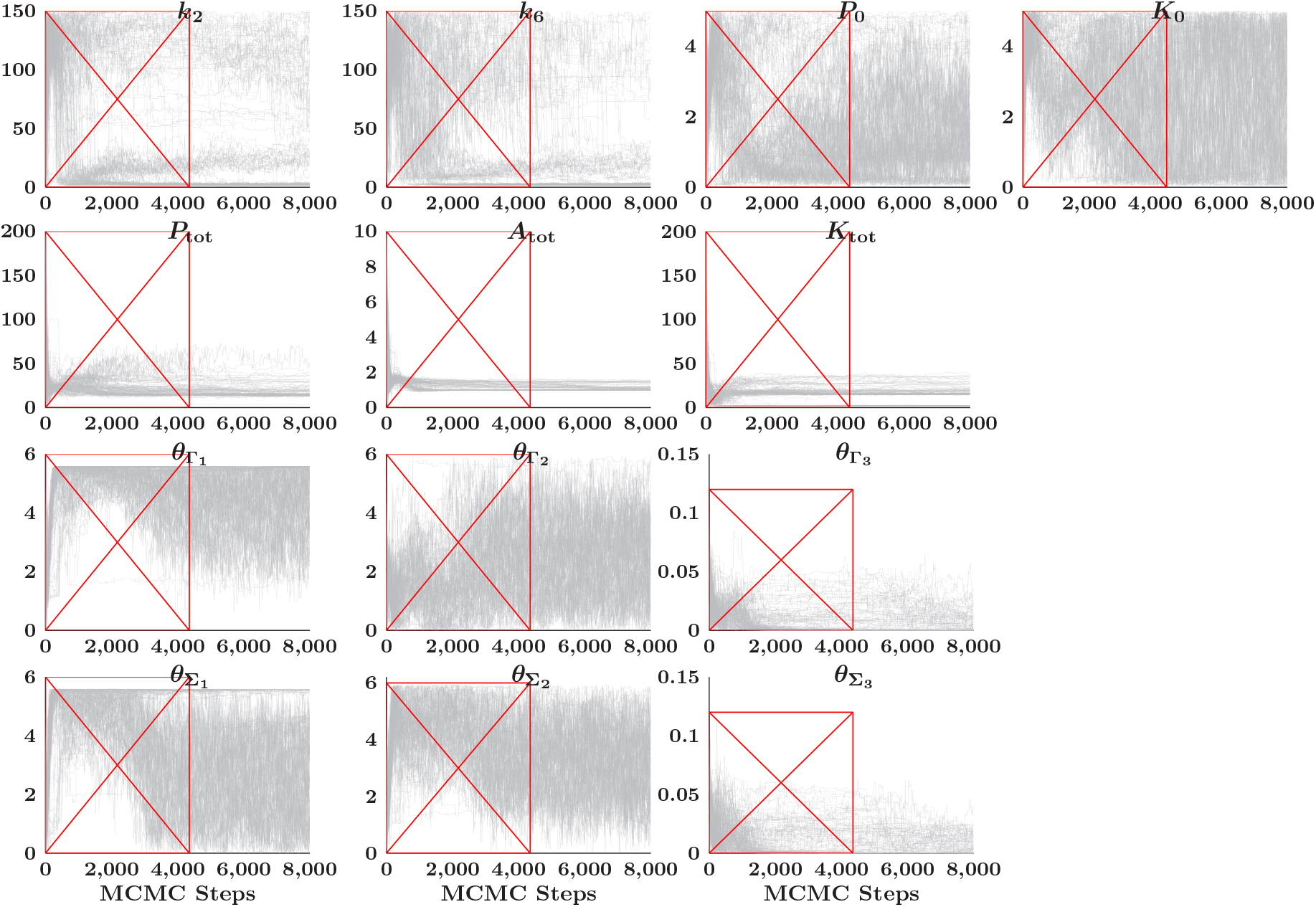
Markov chains for the synaptic plasticity model parameters, Fig 8. The red boxes indicate samples discarded as burn-in.

## Notes

https://github.com/RangamaniLabUCSD/CIUKF-MCMC

### Competing Interest Statement

The authors have declared no competing interest.

## References

[1] Bhalla US, Iyengar R. Robustness of the bistable behavior of a biological signaling feedback loop. Chaos: An Interdisciplinary Journal of Nonlinear Science. 2001;11(1):221–226.

[2] Eungdamrong NJ, Iyengar R. Computational approaches for modeling regulatory cellular networks. Trends in Cell Biology. 2004;14(12):661–669.

[3] Lipshtat A, Jayaraman G, He JC, Iyengar R. Design of versatile biochemical switches that respond to amplitude, duration, and spatial cues. Proceedings of the National Academy of Sciences. 2010;107(3):1247–1252.

[4] Ma’ayan A, Blitzer RD, Iyengar R. Toward predictive models of mammalian cells. Annu Rev Biophys Biomol Struct. 2005;34:319–349.

[5] Song M, Li D, Makaryan SZ, Finley SD. Quantitative modeling to understand cell signaling in the tumor microenvironment. Current Opinion in Systems Biology. 2021;27:100345.

[6] Chakraborty AK. A perspective on the role of computational models in immunology. Annu Rev Immunol. 2017;35:403–439.

[7] Haney S, Bardwell L, Nie Q. Ultrasensitive responses and specificity in cell signaling. BMC systems biology. 2010;4(1):1–14.

[8] Qiao L, Zhao W, Tang C, Nie Q, Zhang L. Network topologies that can achieve dual function of adaptation and noise attenuation. Cell systems. 2019;9(3):271–285.

[9] Wolkenhauer O, Wellstead P, Cho KH, Rangamani P, Iyengar R. Modelling cellular signalling systems. Essays in Biochemistry. 2008;45:83–94.

[10] Zi Z. A tutorial on mathematical modeling of biological signaling pathways. Computational Modeling of Signaling Networks. 2012; p. 41–51.

[11] Keener J, Sneyd J. Mathematical physiology: II: Systems physiology. Springer; 2009.

[12] Mitra ED, Hlavacek WS. Parameter estimation and uncertainty quantification for systems biology models. Curr Opin Syst Biol. 2019;18:9–18.

[13] Babtie AC, Stumpf MPH. How to deal with parameters for whole-cell modelling. J R Soc Interface. 2017;14(133).

[14] Yazdani A, Lu L, Raissi M, Karniadakis GE. Systems biology informed deep learning for inferring parameters and hidden dynamics. PLoS Comput Biol. 2020;16(11):e1007575.

[15] Geris L, Gomez-Cabrero D, editors. Uncertainty in biology: A computational modeling approach. Springer, Cham; 2016.

[16] Raue A, Schilling M, Bachmann J, Matteson A, Schelker M, Kaschek D, et al. Lessons learned from quantitative dynamical modeling in systems biology. PLoS One. 2013;8(9):e74335.

[17] Ashyraliyev M, Fomekong-Nanfack Y, Kaandorp JA, Blom JG. Systems biology: parameter estimation for biochemical models. FEBS J. 2009;276(4):886–902.

[18] Valderrama-Bahamóndez GI, Fröhlich H. MCMC techniques for parameter estimation of ODE based models in systems biology. Frontiers in Applied Mathematics and Statistics. 2019;5:55.

[19] Smith RC. Uncertainty quantification: Theory, implementation, and applications. vol. 12. SIAM; 2013.

[20] Oden JT, Babuška I, Faghihi D. Predictive computational science: Computer predictions in the presence of uncertainty. Encyclopedia of Computational Mechanics Second Edition. 2017; p. 1–26.

[21] Stuart AM. Inverse problems: A Bayesian perspective. Acta numerica. 2010;19:451–559.

[22] Sullivan TJ. Introduction to uncertainty quantification. vol. 63. Springer; 2015.

[23] Kennedy MC, O’Hagan A. Bayesian calibration of computer models. Journal of the Royal Statistical Society: Series B (Statistical Methodology). 2001;63(3):425–464.

[24] Babtie AC, Kirk P, Stumpf MPH. Topological sensitivity analysis for systems biology. Proc Natl Acad Sci U S A. 2014;111(52):18507–18512.

[25] Morrison RE, Oliver TA, Moser RD. Representing model inadequacy: A stochastic operator approach. SIAM/ASA Journal on Uncertainty Quantification. 2018;6(2):457–496.

[26] Galioto N, Gorodetsky AA. Bayesian system ID: Optimal management of parameter, model, and measurement uncertainty. Nonlinear Dyn. 2020;102(1):241–267.

[27] Gelman A, Carlin J, Stern H, Dunson D, Vehtari A, Rubin D. Bayesian data analysis. 3rd ed. CRC Press; 2013.

[28] Gutenkunst RN, Waterfall JJ, Casey FP, Brown KS, Myers CR, Sethna JP. Universally sloppy parameter sensitivities in systems biology models. PLoS Comput Biol. 2007;3(10):1871–1878.

[29] Wieland FG, Hauber AL, Rosenblatt M, Tönsing C, Timmer J. On structural and practical identifiability. Current Opinion in Systems Biology. 2021;25:60–69.

[30] Lillacci G, Khammash M. Parameter estimation and model selection in computational biology. PLoS Comput Biol. 2010;6(3):e1000696.

[31] Schmiester L, Schälte Y, Bergmann FT, Camba T, Dudkin E, Egert J, et al. PEtab–interoperable specification of parameter estimation problems in systems biology. PLoS computational biology. 2021;17(1):e1008646.

[32] Kent E, Neumann S, Kummer U, Mendes P. What can we learn from global sensitivity analysis of biochemical systems? PLoS One.2013;8(11):e79244.

[33] Anstett-Collin F, Denis-Vidal L, Millérioux G. A priori identifiability: An overview on definitions and approaches. Annu Rev Control. 2020;50:139–149.

[34] Erguler K, Stumpf MPH. Practical limits for reverse engineering of dynamical systems: A statistical analysis of sensitivity and parameter inferability in systems biology models. Mol Biosyst. 2011;7(5):1593–1602.

[35] Raue A, Kreutz C, Maiwald T, Bachmann J, Schilling M, Klingmüller U, et al. Structural and practical identifiability analysis of partially observed dynamical models by exploiting the profile likelihood. Bioinformatics. 2009;25(15):1923–1929.

[36] Hong H, Ovchinnikov A, Pogudin G, Yap C. Global identifiability of differential models. Commun Pure Appl Math. 2020;73(9):1831–1879.

[37] Saltelli A, Ratto M, Andres T, Campolongo F, Cariboni J, Gatelli D, et al. Global sensitivity analysis. The primer. Wiley; 2008.

[38] Varma A, Morbidelli M, Wu H. Parametric sensitivity in chemical systems. Varma A, editor. Cambridge Series in Chemical Engineering. Cambridge University Press; 1999.

[39] Marino S, Hogue IB, Ray CJ, Kirschner DE. A methodology for performing global uncertainty and sensitivity analysis in systems biology. J Theor Biol. 2008;254(1):178–196.

[40] Mortlock RD, Georgia SK, Finley SD. Dynamic regulation of JAK-STAT signaling through the prolactin receptor predicted by computational modeling. Cell Mol Bioeng. 2021;14(1):15–30.

[41] Kay SM. Fundamentals of statistical signal processing: estimation theory. Prentice-Hall, Inc.; 1993.

[42] Moon TK, Stirling WC. Mathematical methods and algorithms for signal processing. 621.39: 51 MON. Prentice Hall; 2000.

[43] Liepe J, Kirk P, Filippi S, Toni CP Tinaand Barnes, Stumpf MPH. A framework for parameter estimation and model selection from experimental data in systems biology using approximate Bayesian computation. Nat Protoc. 2014;9(2):439–456.

[44] Toni T, Welch D, Strelkowa N, Ipsen A, Stumpf MPH. Approximate Bayesian computation scheme for parameter inference and model selection in dynamical systems. J R Soc Interface. 2009;6(31):187–202.

[45] Golightly A, Wilkinson DJ. Bayesian parameter inference for stochastic biochemical network models using particle Markov chain Monte Carlo. Interface Focus. 2011;1(6):807–820.

[46] Ghasemi O, Lindsey ML, Yang T, Nguyen N, Huang Y, Jin YF. Bayesian parameter estimation for nonlinear modelling of biological pathways. BMC Syst Biol. 2011;5 Suppl 3:S9.

[47] Bianconi F, Antonini C, Tomassoni L, Valigi P. Application of conditional robust calibration to ordinary differential equations models in computational systems biology: A comparison of two sampling strategies. IET Syst Biol. 2020;14(3):107–119.

[48] Wilkinson DJ. Bayesian methods in bioinformatics and computational systems biology. Brief Bioinform. 2007;8(2):109–116.

[49] Klinke DJ 2nd. An empirical Bayesian approach for model-based inference of cellular signaling networks. BMC Bioinformatics. 2009;10:371.

[50] Renardy M, Yi TM, Xiu D, Chou CS. Parameter uncertainty quantification using surrogate models applied to a spatial model of yeast mating polarization. PLoS Comput Biol. 2018;14(5):e1006181.

[51] Erazo K, Nagarajaiah S. An offline approach for output-only Bayesian identification of stochastic nonlinear systems using unscented Kalman filtering. J Sound Vib. 2017;397:222–240.

[52] Hong H, Ovchinnikov A, Pogudin G, Yap C. SIAN: Software for structural identifiability analysis of ODE models. Bioinformatics. 2019;35(16):2873–2874.

[53] Sobol IM. Global sensitivity indices for nonlinear mathematical models and their Monte Carlo estimates. Math Comput Simul. 2001;55:271–280.

[54] Teixeira B, Torres LAB, Aguirre LA, Bernstein DS. Unscented filtering for interval-constrained nonlinear systems. In: Proceedings of the 47th IEEE Conference on Decision and Control, CDC 2008, December 9-11, 2008, Cancún, México. Institute of Electrical and Electronics Engineers; 2008. p. 5116–5121.

[55] Villaverde AF, Evans ND, Chappell MJ, Banga JR. Input-Dependent structural identifiability of nonlinear systems. IEEE Control Systems Letters. 2019;3(2):272–277.

[56] Nguyen LK, Degasperi A, Cotter P, Kholodenko BN. DYVIPAC: An integrated analysis and visualisation framework to probe multi-dimensional biological networks. Sci Rep. 2015;5:12569.

[57] Pi HJ, Lisman JE. Coupled phosphatase and kinase switches produce the tristability required for long-term potentiation and long-term depression. J Neurosci. 2008;28(49):13132–13138.

[58] Haario H, Laine M, Mira A, Saksman E. DRAM: Efficient adaptive MCMC. Stat Comput. 2006;16(4):339–354.

[59] Goodman J, Weare J. Ensemble samplers with affine invariance. Communications in Applied Mathematics and Computational Science. 2010;5(1):65– 80.

[60] Norton J, Walter E, Pronzato L. Identification of parametric models from experimental data. Communications and Control Engineering. Springer London; 2010.

[61] Ljung L, Glad T. On global identifiability for arbitrary model parametrizations. Automatica. 1994;30(2):265–276.

[62] Villaverde AF. Observability and structural identifiability of nonlinear biological systems. Complexity. 2019;2019.

[63] Bezanson J, Edelman A, Karpinski S, Shah VB. Julia: A fresh approach to numerical computing. SIAM review. 2017;59(1):65–98.

[64] Ilia I, Ovchinnikov A, Pogudin G. SIAN.jl-Implementation of SIAN in Julia; 2022. https://github.com/alexeyovchinnikov/SIAN-Julia.

[65] Marelli S, Sudret B. UQLab: A framework for uncertainty quantification in Matlab. In: Vulnerability, uncertainty, and risk: quantification, mitigation, and management; 2014. p. 2554–2563.

[66] Marelli S, Lamas C, Konakli K, Mylonas C, Wiederkehr P, Sudret B. UQLab user manual – Sensitivity analysis. Chair of Risk, Safety and Uncertainty Quantification, ETH Zurich, Switzerland; 2022.

[67] Rackauckas C, Nie Q. Differentialequations.jl–a performant and feature-rich ecosystem for solving differential equations in julia. Journal of Open Research Software. 2017;5(1).

[68] Särkkä S. Bayesian filtering and smoothing. 3. Cambridge University Press; 2013.

[69] Julier SJ, Uhlmann JK. Unscented filtering and nonlinear estimation. Proc IEEE. 2004;92(3):401–422.

[70] Vachhani P, Narasimhan S, Rengaswamy R. Robust and reliable estimation via Unscented Recursive Nonlinear Dynamic Data Reconciliation. J Process Control. 2006;16(10):1075–1086.

[71] Simon D. Kalman filtering with state constraints: a survey of linear and nonlinear algorithms. IET Control Theory Appl. 2010;4(8):1303–1318.

[72] Julier SJ, Uhlmann JK. New extension of the Kalman filter to nonlinear systems. In: Signal processing, sensor fusion, and target recognition VI. vol. 3068. International Society for Optics and Photonics; 1997. p. 182–193.

[73] Tsigkinopoulou A, Hawari A, Uttley M, Breitling R. Defining informative priors for ensemble modeling in systems biology. Nat Protoc. 2018;13(11):2643–2663.

[74] Gelman A. Prior distributions for variance parameters in hierarchical models (comment on article by Browne and Draper). Bayesian Analysis. 2006;1(3):515–534.

[75] Gelman A, Roberts G, Gilks W. Efficient Metropolis jumping hules. Bayesian statistics. 1996;.

[76] Owen AB. Monte Carlo theory, methods and examples; 2013.

[77] Sokal A. Monte Carlo methods in statistical mechanics: Foundations and new algorithms. In: Functional integration. Springer; 1997. p. 131–192.

[78] Metropolis N, Rosenbluth AW, Rosenbluth MN, Teller AH, Teller E. Equation of state calculations by fast computing machines. The journal of chemical physics. 1953;21(6):1087–1092.

[79] Hastings WK. Monte Carlo sampling methods using Markov chains and their applications. Biometrika. 1970;57(1):97–109.

[80] Tierney L. Markov chains for exploring posterior distributions. the Annals of Statistics. 1994; p. 1701–1728.

[81] Haario H, Saksman E, Tamminen J. An adaptive Metropolis algorithm. Bernoulli. 2001; p. 223–242.

[82] Wagner PR, Nagel J, Marelli S, Sudret B. UQLab user manual–Bayesian inversion for model calibration andvalidation. Chair of Risk, Safety and Uncertainty Quantification, ETH Zurich, Switzerland; 2022.

[83] Wolff U. Monte Carlo errors with less errors. Comput Phys Commun. 2004;156(2):143–153.

[84] Mišković L, Hatzimanikatis V. Modeling of uncertainties in biochemical reactions. Biotechnol Bioeng. 2011;108(2):413–423.

[85] Bowman AW, Azzalini A. Applied smoothing techniques for data analysis: The kernel approach with S-Plus illustrations. vol. 18. OUP Oxford; 1997.

[86] Shankaran H, Wiley HS. Oscillatory dynamics of the extracellular signal-regulated kinase pathway. Curr Opin Genet Dev. 2010;20(6):650–655.

[87] Shaul YD, Seger R. The MEK/ERK cascade: From signaling specificity to diverse functions. Biochimica et Biophysica Acta (BBA)-Molecular Cell Research. 2007;1773(8):1213–1226.

[88] Vera J, Rath O, Balsa-Canto E, Banga JR, Kolch W, Wolkenhauer O. Investigating dynamics of inhibitory and feedback loops in ERK signalling using power-law models. Mol Biosyst. 2010;6(11):2174–2191.

[89] Kholodenko BN. Negative feedback and ultrasensitivity can bring about oscillations in the mitogen-activated protein kinase cascades. Eur J Biochem. 2000;267(6):1583–1588.

[90] Kholodenko BN, Hancock JF, Kolch W. Signalling ballet in space and time. Nat Rev Mol Cell Biol. 2010;11(6):414–426.

[91] Äijö T, Bonneau R. Biophysically motivated regulatory network inference: Progress and prospects. Hum Hered. 2016;81(2):62–77.

[92] Casadiego J, Nitzan M, Hallerberg S, Timme M. Model-free inference of direct network interactions from nonlinear collective dynamics. Nat Commun. 2017;8(1):2192.

[93] Schmidt M, Lipson H. Distilling free-form natural laws from experimental data. Science. 2009;324(5923):81–85.

[94] Brunton SL, Proctor JL, Kutz JN. Discovering governing equations from data by sparse identification of nonlinear dynamical systems. Proc Natl Acad Sci U S A. 2016;113(15):3932–3937.

[95] Mangan NM, Brunton SL, Proctor JL, Kutz JN. Inferring biological networks by sparse identification of nonlinear dynamics. IEEE Transactions on Molecular, Biological and Multi-Scale Communications. 2016;2(1):52–63.

[96] Hoffmann M, Fröhner C, Noé F. Reactive SINDy: Discovering governing reactions from concentration data. J Chem Phys. 2019;150(2):025101.

[97] Kaheman K, Kutz JN, Brunton SL. SINDy-PI: a robust algorithm for parallel implicit sparse identification of nonlinear dynamics. Proc Math Phys Eng Sci. 2020;476(2242):20200279.

[98] Hirsh SM, Barajas-Solano DA, Kutz JN. Sparsifying priors for Bayesian uncertainty quantification in model discovery. arXiv preprint arXiv:210702107. 2021;.

[99] Foreman-Mackey D, Hogg DW, Lang D, Goodman J. emcee: The MCMC Hammer. PASP. 2013;125(925):306.

[100] Blei DM, Kucukelbir A, McAuliffe JD. Variational inference: A review for statisticians. J Am Stat Assoc. 2017;112(518):859–877.

[101] Bardsley JM, Solonen A, Haario H, Laine M. Randomize-then-optimize: A method for sampling from posterior distributions in nonlinear inverse problems. SIAM Journal on Scientific Computing. 2014;36(4):A1895–A1910.

[102] Gasca-Aragon H. Data combination from multiple sources under measurement error. University of Massachusetts Amherst; 2012.

[103] Marin-Martinez F, Sánchez-Meca J. Weighting by inverse variance or by sample size in randomeffects meta-analysis. Educational and Psychological Measurement. 2010;70(1):56–73.

[104] Clemen RT, Winkler RL. Combining probability distributions from experts in risk analysis. Risk analysis. 1999;19(2):187–203.

